# Zebrafish fin regeneration requires generic and regeneration-specific responses of osteoblasts to trauma

**DOI:** 10.1101/2022.02.22.481466

**Authors:** Ivonne Sehring, Melanie Haffner-Luntzer, Anita Ignatius, Markus Huber-Lang, Gilbert Weidinger

**Affiliations:** Institute of Biochemistry and Molecular Biology, Ulm University, Albert-Einstein-Allee 11, 89081 Ulm, Germany; Institute of Orthopaedic Research and Biomechanics, Ulm Medical Center, University Hospital Ulm, Helmholtzstrasse 14, 89081 Ulm, Germany; Institute of Clinical and Experimental Trauma-Immunology (ITI), University Hospital Ulm, Helmholtzstrasse 8/1, 89081 Ulm, Germany

**Keywords:** zebrafish, regeneration, bone, osteoblast, dedifferentiation, migration, complement system, C5a, C5aR1, C3a, NF-κB, NF-kappaB, retinoic acid, actomyosin, microtubuli, fin, trauma

## Abstract

Successful regeneration requires the coordinated execution of multiple cellular responses to injury. In amputated zebrafish fins, mature osteoblasts dedifferentiate, migrate towards the injury and form proliferative osteogenic blastema cells. We show that osteoblast migration is preceded by cell elongation and alignment along the proximodistal axis, which require actomyosin, but not microtubule turnover. Surprisingly, osteoblast dedifferentiation and migration can be uncoupled. Using pharmacological and genetic interventions, we found that NF-κB and retinoic acid signalling regulate dedifferentiation without affecting migration, while the complement system and actomyosin dynamics are required for migration but not dedifferentiation. Furthermore, by removing bone at two locations within a fin ray, we established a trauma model containing two injury sites. We found that osteoblasts dedifferentiate at and migrate towards both sites, while accumulation of osteogenic progenitor cells and regenerative bone formation only occur at the distal-facing injury. Together, these data indicate that osteoblast dedifferentiation and migration represent generic injury responses that are differentially regulated and can occur independently of each other and of regenerative growth. Successful bone regeneration appears to require the coordinated execution of generic and regeneration-specific responses of osteoblast to trauma.

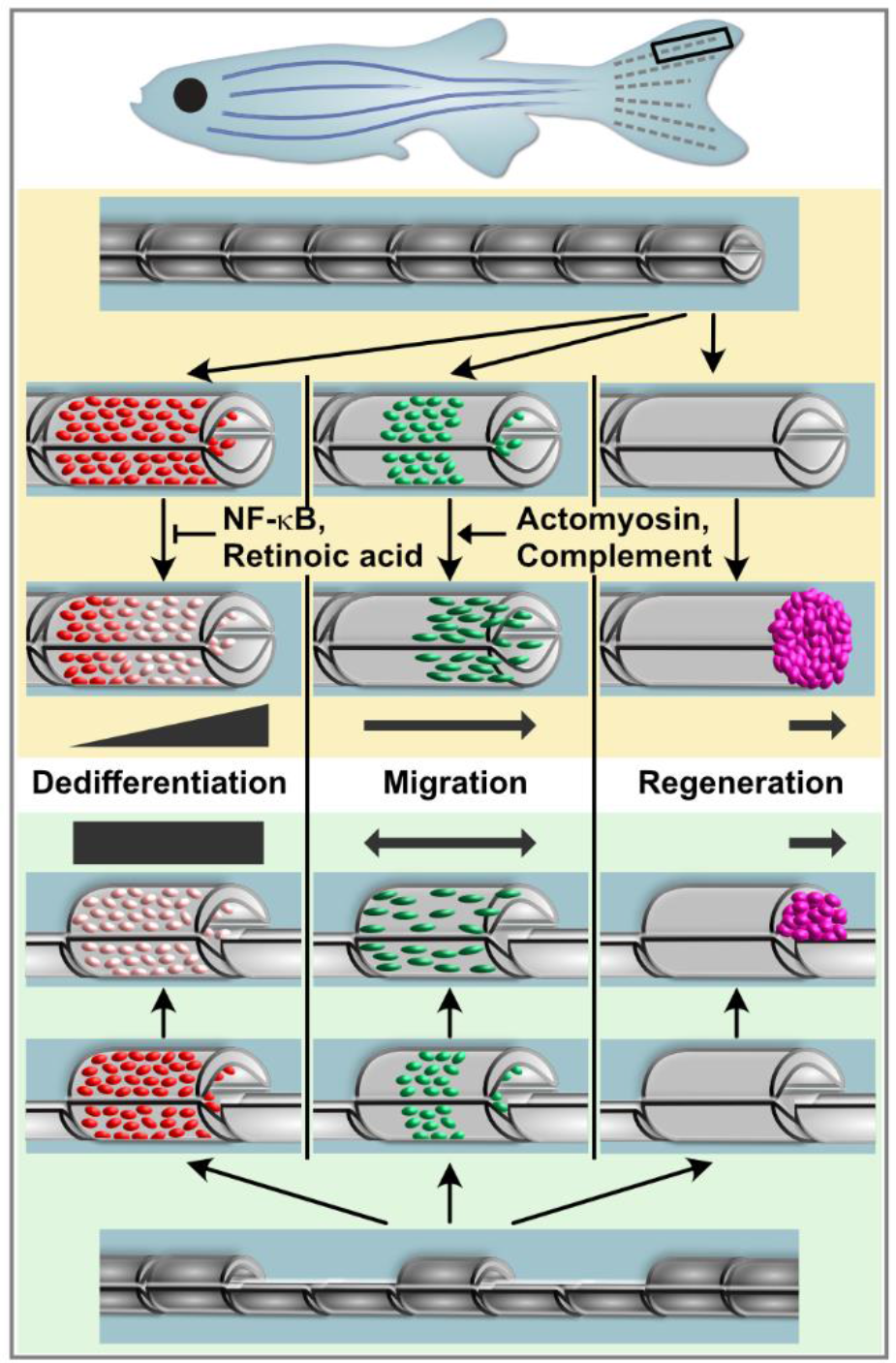

## Introduction

For humans and other mammals, the traumatic loss of a limb represents an irreversible and permanent defect, as lost appendages cannot be restored. In contrast, teleost fish and urodele amphibians are able to fully regenerate limbs / fins. Thus, the zebrafish caudal fin has become a popular model to study the restoration of bone tissue (Gemberling *et al*., 2013; Pfefferli and Jazwinska, 2015). The fin skeleton consists of exoskeletal elements, the fin rays or lepidotrichia, which stretch across the whole fin, and endochondral parts close to the body. Rays are segmented with flexible joints, and each segment consists of two concave hemirays which are lined by a single layer of osteoblasts on the inner and outer surface (Figure 1A, B). The fin rays are key regenerative units. After amputation, within one day a wound epidermis covers the injured tissue. Next, a blastema forms atop of each ray, a hallmark for epimorphic regeneration. Blastema cells are responsible for proliferation and forming new structures by differentiating into mature cells. Within hours after amputation, osteoblasts close to the amputation plane dedifferentiate, that is they downregulate the expression of the mature osteoblast marker *bglap* and upregulate pre-osteoblast markers like *runx2* (Knopf *et al*., 2011; Sousa *et al*., 2011; Stewart and Stankunas, 2012). Furthermore, osteoblasts relocate towards the amputation plane and beyond to contribute to the blastema (Knopf *et al*., 2011; Geurtzen *et al*., 2014). In the regenerate, these dedifferentiated osteoblasts remain lineage-restricted and re-differentiate to osteoblasts (Knopf *et al*., 2011; Stewart and Stankunas, 2012). While both osteoblast dedifferentiation and migration occur in response to fin amputation and bone fractures in zebrafish (Knopf *et al*., 2011; Geurtzen *et al*., 2014), the interrelation of these processes is not understood; specifically, whether dedifferentiation is a requirement for migration is not known.

**Figure 1.**
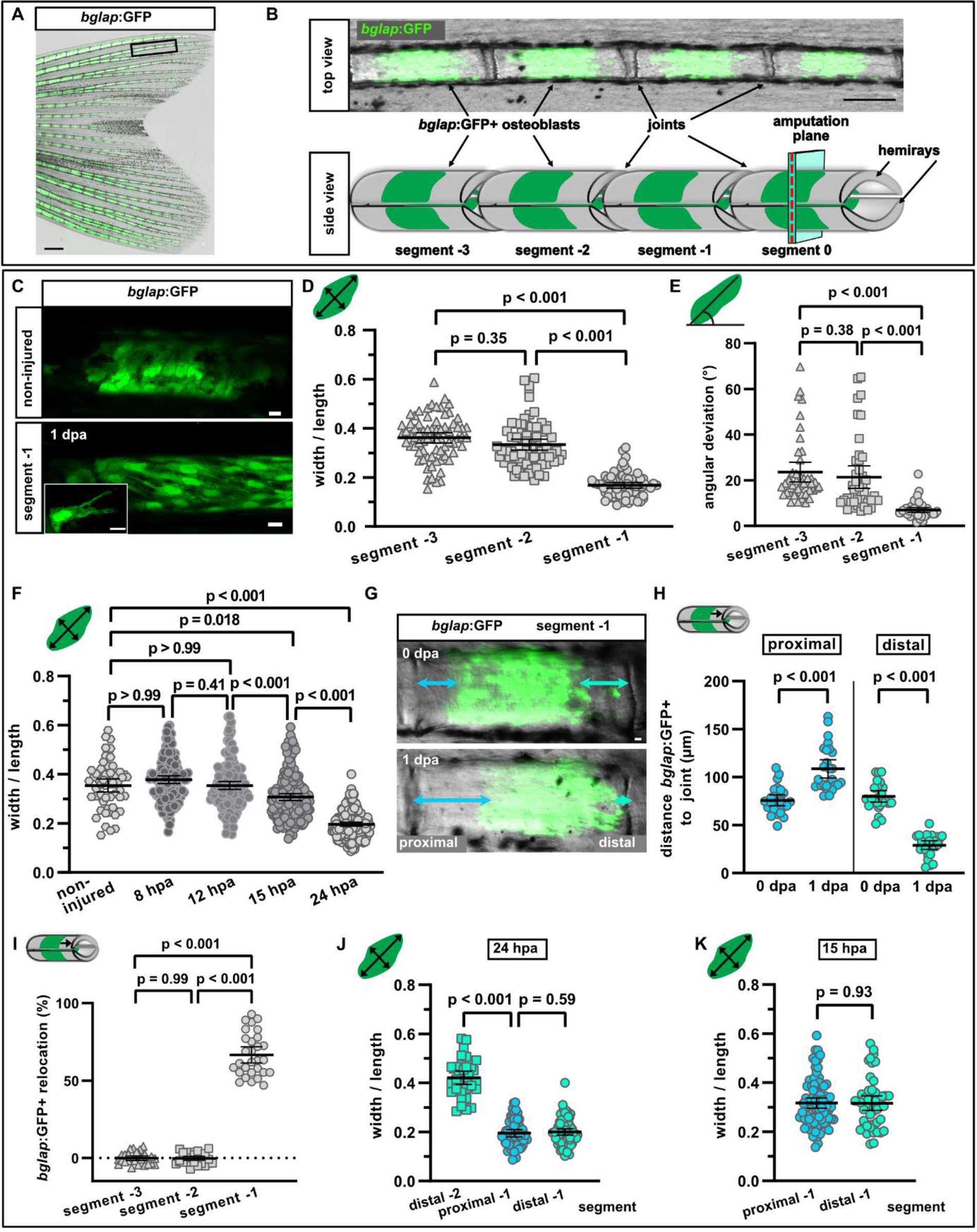
Osteoblast morphology change and migration. A) Adult zebrafish fin expressing *bglap*:GFP in mature osteoblasts. Scale bar, 500 µm. B) In non-injured fins, *bglap*:GFP is expressed in a subset of osteoblasts in the centre of each segment, and absent around the joints. Scale bar, 100 µm. C) *bglap*:GFP+ osteoblast morphology in a non-injured fin (upper panel) and in segment −1 at 1 dpa (lower panel). Scale bar, 10 µm. D) Quantification of *bglap*:GFP+ osteoblast roundness as width / length ratio at 1 dpa. Osteoblasts are elongated only in segment −1. n (fins) = 5, n (rays) = 5, n (cells) = 72. Error bars represent 95% CI. Kruskal-Wallis test. E) Quantification of *bglap*:GFP+ osteoblast orientation as angular deviation from the proximodistal axis at 1 dpa. In segment −1, osteoblasts are aligned along the axis. n (fins) = 5, n (rays) = 5, n (cells) = 44. Error bars represent 95% CI. Kruskal-Wallis test. F) Osteoblast roundness at various time points. Elongation can first be detected at 15 hpa. n (rays) = 5 (non-injured), 10 (8 hpa), 8 (12 hpa), 12 (15 hpa), 11 (24 hpa); n (cells) = 54 (non-injured), 134 (8 hpa), 151 (12 hpa), 176 (15 hpa), 176 (24 hpa). Error bars represent 95% CI. Kruskal- Wallis test. G) Live imaging of *bglap*:GFP+ cells in segment −1 at 0 and 1 dpa showing the relocation towards the distal joint. Distal to the right. Scale bar, 10 µm. H) Quantification of the distance between the bulk of *bglap*:GFP+ cells and joints in segment −1 at 0 and 1 dpa. At 1 dpa, the proximal distance is increased, while the distal distance is reduced. n (fins) = 26. Error bars represent 95% CI. Unpaired t-test. I) Quantification of *bglap*:GFP+ bulk migration in segment −3, −2 and −1 at 1 dpa. 100% indicates full crossing of the distance to the respective joint. Osteoblasts migrate distally in segment −1. n (fins) = 19, n (rays) = 29. Error bars represent 95% CI. Mann Whitney test. J) Osteoblast roundness in distal and proximal parts of a segment (40% of total segment length from the respective end) at 24 hpa. No difference within segment −1 can be detected. n (fins) = 5, n (rays) = 5, n (cells) = 34 (segment −2), 55 (proximal segment −1), 78 (distal segment −1). Error bars represent 95% CI. Mann-Whitney test. K) Osteoblast roundness in distal and proximal parts of segment −1 at 15 hpa. No difference within segment −1 can be detected. n (fins) = 6, n (rays) = 15, n (cells) = 82 (proximal), 49 (distal). Error bars represent 95% CI. Mann-Whitney test. The observed relative difference is 0.3%, the calculated smallest significant difference 13% and thus smaller than what we observe between segment −2 and segment −1 (Figure 1J, 54%).

Cell migration involves the formation of protrusive membrane structures such as actin-rich protrusions, pseudopodia, and blebs (Mierke, 2015). Their formation is regulated by extrusive and contractile forces of the cytoskeleton, which is mainly composed of actin microfilaments, microtubules, and intermediate filaments. The dynamic actomyosin network is important for formation and retraction of membrane protrusions, while microtubuli play a role in organizing the polarization of migrating cells (Petrie, Doyle and Yamada, 2009). Bone tissue is permanently turned over by bone remodelling, a process of alternating bone resorption and bone formation (Kular *et al*., 2012). For bone formation, osteoblasts migrate to the resorbed sites. Similarly, during mammalian fracture healing, osteoblasts are recruited to the site of injury (Thiel *et al*., 2018). Hence, for efficient recruitment, osteoblasts must sense and respond to specific signals. Several factors have been shown to act as chemoattractant for osteoblasts *in vitro*, but few have been confirmed to play a role in vivo (Dirckx, Van Hul and Maes, 2013; Thiel *et al*., 2018). One candidate guidance cue for osteoblasts is the complement system, which represents the major fluid phase part of innate immunity, and is activated immediately after trauma. It consists of more than 50 proteins, including serial proteases, whose activation leads to the formation of the anaphylatoxin peptides C3a and C5a, generated from the precursors C3 and C5, respectively (Thorgersen *et al*., 2019). It has been shown that C3 and C5 mRNA are expressed by human osteoblasts, and that they can cleave native C5 into C5a (Ignatius, Schoengraf, *et al*., 2011). During bone fracture healing in mammals, the receptor for C5a (C5aR) is expressed by osteoblasts, and C5a can act as chemoattractant for osteoblasts *in vitro* (Ignatius, *et al*., 2011). In C5-deficient mice, bone repair after fracture is severely impaired (Ehrnthaller *et al*., 2013), however, the underlying mechanism of action is not yet understood.

In this study, we analysed injury-induced osteoblast migration *in vivo* in the regenerating zebrafish fin. We show that osteoblast cell shape changes and migration depend on a dynamic actomyosin cytoskeleton, but not on microtubuli turnover. The complement factors C3a and C5a are required for osteoblast migration. Using genetic and pharmacological manipulation of NF-κB, retinoic acid and complement signalling, we found that dedifferentiation and migration can be uncoupled and are independently regulated, suggesting that dedifferentiation is not a prerequisite for migration. Furthermore, we established a novel injury model in which an internal bone defect within fin rays allows us to study osteoblast behaviours at proximally and distally facing injuries. Intriguingly, osteoblast migration and dedifferentiation occur at both injury sites, yet only at the distal injury a pre-osteoblast population forms and only here regenerative growth commences. We conclude that osteoblast migration and dedifferentiation represent generic injury responses in zebrafish, and that successful bone regeneration depends on additional, regeneration-specific events.

## Results

### Osteoblasts elongate, align along the proximodistal axis and migrate towards the amputation plane

To observe osteoblast behaviour at single cell resolution in live fish after fin amputation, we generated a double transgenic line expressing *bglap*:GFP (Knopf *et al*., 2011) and *entpd5*:kaede (Huitema *et al*., 2012) in mature osteoblasts. Partial photoconversion of kaedeGreen into kaedeRed resulted in a different colouring for each cell which allows single cell tracking. Live imaging revealed the formation of long protrusions, lasting for at least 2 h and extending towards the amputation site, and the directed movement of cell bodies relative to their surroundings (Video 1, Figure 1 – figure supplement 1A). To quantify osteoblast behaviour, we used the *bglap*:GFP transgenic line. In adult non-injured fin rays, *bglap*:GFP is expressed in a subset of mature osteoblasts in a segmented pattern (Figure 1A). This pattern results from the localization of *bglap*:GFP+ cells to the centre of every ray segment, while they are excluded from the flexible segment boundaries (Figure 1B). Migrating cells typically possess an identifiable cell front and a cell rear along an axis approximately aligned with the direction of locomotion. Therefore, we analysed if a change in osteoblast shape can be observed after amputation. We use the following terminology to describe the subsequent experiments: segment 0 is the fin ray segment through which we amputate (at 50% of its length), segment − 1 is the segment located proximally to segment 0, and segments −2 and segment −3 those located even further proximally. Note that only segment 0 is mechanically affected by the amputation injury. We found that *bglap*:GFP+ osteoblasts drastically changed their shape after amputation. In a mature, non-injured segment, osteoblasts were roundish, and they retained this morphology after amputation in segment −3 and segment −2 at 1 day post amputation (dpa), as determined by a width / length ratio of ∼ 0.4 (Figure 1C, D, Figure 1 – figure supplement 1B). In contrast, at 1 dpa osteoblasts in segment −1 displayed an elongated shape (width / length ratio ∼ 0.2) and they presented long extensions (Figure 1C, D). The elongation of osteoblasts occurred in alignment with the proximodistal axis of the fin, as evident by an angle reduction between this axis and the long axis of the osteoblasts (Figure 1E). Orientation of osteoblasts along the proximodistal axis could first be detected at 12 hours post amputation (hpa) (Figure 1 – figure supplement 1C), while elongation was first observed at 15 hpa (Figure 1F). We interpret the elongation and orientation of osteoblasts along the proximodistal axis and the formation of long-lived protrusions along this axis as evidence for directed active migration behaviour towards the injury.

The restriction of *bglap*:GFP+ cells to the segment centre results in a zone devoid of *bglap*:GFP+ cells at both ends of a segment and its invasion by GFP+ cells can be used as a read-out for bulk migration of osteoblasts. Within 1 dpa, the distance between the GFP+ cells and the distal joint was reduced in segment −1, while at the proximal joint, the distance increased (Figure 1G, H). Therefore, osteoblasts did not spread out across the segment, but were relocated directionally towards the amputation plane. We also analysed if osteoblasts in more proximal segments (further away from the amputation plane) migrated distally and quantified their relocation. 100% relocation indicates that the distal front of the GFP+ bulk of cells has reached the respective distal joint. At 1 dpi, osteoblasts in segment −1 migrated distally, while no migration could be detected in segments −3 and −2 (Figure 1I).

Fins grow by the addition of new segments distally, and in the distal-most, youngest segment, *bglap*:GFP is not expressed (Knopf *et al*., 2011). Thus, during ontogenetic growth and in regenerating fins at later stages where osteoblasts re-differentiate, increasing numbers of GFP+ cells can be detected in more proximally located, older segments, reflecting the progressive differentiation of osteoblasts with time (Figure 1 – figure supplement 1D). In segments that start to upregulate *bglap* expression, all *bglap*:GFP+ cells appear in the centre of the segments; we could not observe GFP+ cells within joints (Figure 1 – figure supplement 1D). This suggests that the mature osteoblast population in a segment is formed via differentiation of osteoblasts at the position within the segment where they were formed during segment addition, and not via migration of differentiated, *bglap*:GFP+ osteoblasts from older segments into less mature segments. In contrast, within 2 days after fin amputation, *bglap*:GFP+ cells appeared within the joint between segment −1 and segment 0 (yellow arrowhead) (Figure 1 – figure supplement 1E), indicating that GFP+ osteoblasts from segment −1 crossed the joint during their migration towards the amputation plane. Thus, migration of mature osteoblasts observed after fin amputation or bone fracture (Geurtzen *et al*., 2014) appears to represent a specific early response to trauma, while ontogenetic or regenerative bone formation does not involve migration of differentiated osteoblasts.

We have previously shown that within 1 dpa, osteoblasts dedifferentiate in a zone extending to 350 µm from the amputation plane (Knopf *et al*., 2011). In the experiments presented here, where we always amputated at the centre of a segment, this 350 µm zone comprises the amputated segment 0 and the adjacent segment −1. We conclude that osteoblasts dedifferentiate in the same region where they change their morphology. Using RNAscope in situ hybridization, we found that expression of *bglap* gradually decreases towards the amputation plane at 1 dpa (Figure 1 – figure supplement 1F). Thus, we wondered if a similar gradient can be observed for cell morphology changes. However, osteoblasts were elongated to the same extent at the proximal and distal end of segment −1 at 24 hpa (Figure 1J). As elongation was first observed at 15 hpa (Figure 1F), we also analysed osteoblasts in proximal and distal regions of segment − 1 at this time point. Yet, no morphological differences in osteoblasts of proximal and distal regions of segment −1 were detected (Figure 1K). We conclude that dedifferentiation and cell shape changes are responses of osteoblasts to injury that occur in the same cells, yet the magnitude of these responses can vary. While cell shape change is a binary response that affects all osteoblasts in the responsive zone equally, dedifferentiation is a graded process. In summary, after amputation, osteoblasts in segment −1 drastically change their shape and orientation and migrate towards the amputation site.

### Actomyosin, but not microtubule dynamics is required for osteoblast cell shape changes and migration

Cell shape is largely determined by the cell cortex, and one of the primary forces behind a cell shape change is the myosin-associated actin cytoskeleton (Chugh and Paluch, 2018). The organization of the actin cytoskeleton is also an important factor determining the characteristics of the motility during cell migration (Svitkina, 2018). Therefore, we next analysed if interference with actomyosin dynamics had an effect on osteoblast cell shape change and migration. To interfere with the treadmilling of actin microfilaments (F-actin), we used cytochalasin D. Cytochalasin D binds to F-actin and disturbs its turnover, ultimately changing F-actin dynamics (Brown and Spudich, 1979, 1981). Drug treatment strongly impaired osteoblast elongation in segment −1 at 1 dpa, while it did not affect osteoblast morphology in segment −2 and segment −3 (Figure 2A). Concomitantly, treatment with cytochalasin D also resulted in reduced alignment of osteoblasts along the proximodistal axis in segment −1, but did not affect the orientation of osteoblasts in segment −2 and segment −3 (Figure 2B). Interference with actin dynamics also significantly reduced the bulk migration of *bglap*:GFP+ cells in segment −1 (Figure 2C). To address another hallmark of a dynamic cytoskeleton, the contractility of the actomyosin network, we injected fish with blebbistatin, an inhibitor of myosin II ATPase (Kovács *et al*., 2004). Drug treatment did not affect osteoblast cell shape in segments −3 and −2, but impaired the elongation and re-orientation of osteoblasts along the proximodistal axis in segment −1 (Figure 2D, E). Concomitantly, bulk osteoblast migration was reduced (Figure 2C). *Bglap*:GFP transgenic fish can also be used as live reporters for osteoblast dedifferentiation, as downregulation of *bglap* expression can be monitored as reduction of GFP intensity at 3 dpa (Knopf *et al*., 2011; Mishra *et al*., 2020). Importantly, osteoblast dedifferentiation was not affected by either cytochalasin D nor blebbistatin treatment (Figure 2F), indicating that the effect on cell shape change and migration is not secondary due to impaired dedifferentiation.

**Figure 2.**
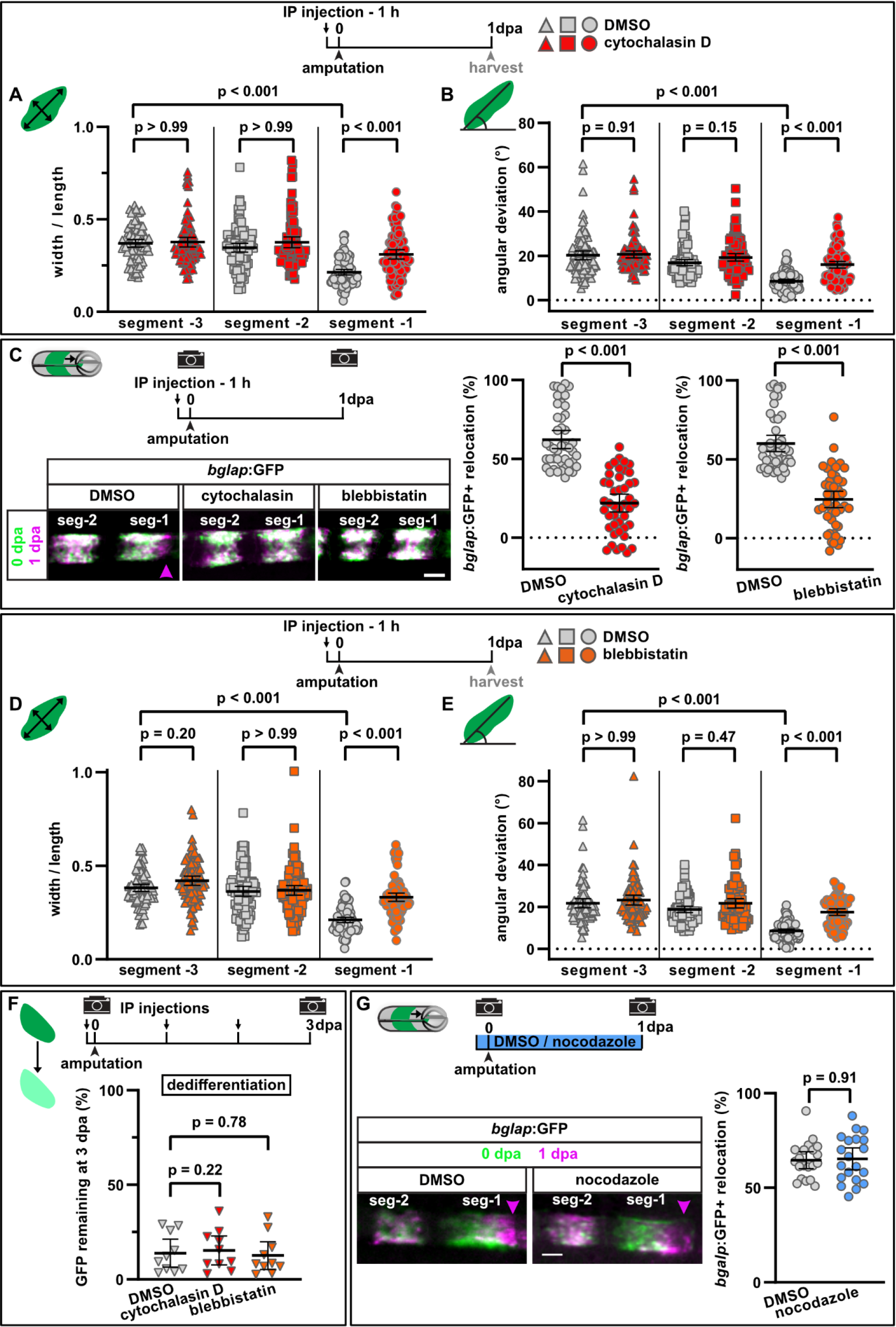
Interference with actomyosin dynamics affects osteoblast cell shape change and migration. A) Osteoblast roundness at 1 dpa. Inhibition of actin dynamics with cytochalasin D does not alter osteoblast cell shape in segments −3 and −2, but cell elongation in segment −1 is inhibited. n (fins) = 15, n (rays) = 15, n (cells) = 87. Error bars represent 95% CI. Kruskal-Wallis test. B) Osteoblast orientation at 1 dpa. Cytochalasin D does not alter osteoblast orientation in segments −3 and −2, but alignment along the proximodistal axis in segment −1 is impaired. N (experiments) = 3, n (fins) = 15, n (rays) = 15, n (cells) = 93. Error bars represent 95% CI. Kruskal-Wallis test. C) Both cytochalasin D and blebbistatin treatment impair bulk osteoblast migration. Images show overlay of 0 dpa (green) and 1 dpa (pink) pictures, with the pink arrowhead indicating relocation of osteoblasts in controls, where no signal overlap is observed at the distal side. Graph: 100% indicates full crossing of the distance to the respective joint at 1 dpa. Cytochalasin D: N (experiments) = 3, n (fins) = 22, n (rays) = 44; blebbistatin: n (fins) = 24, n (rays) = 48, appertaining controls have the same n. Error bars represent 95% CI. Mann-Whitney test. Scale bar, 100 µm. D) Osteoblast roundness at 1 dpa. Inhibition of myosin activity with blebbistatin does not alter osteoblast cell shape in segments −3 and −2, but cell elongation in segment −1 is inhibited. N (experiments) = 3, n (fins) = 15, n (rays) = 15, n (cells) = 89. Error bars represent 95% CI. Kruskal-Wallis test. E) Osteoblast orientation at 1 dpa. Blebbistatin does not alter osteoblast orientation in segments −3 and −2, but alignment along the proximodistal axis in segment −1 is impaired. N (experiments) = 3, n (fins) = 15, n (rays) = 15, n (cells) = 77. Error bars represent 95% CI. Kruskal-Wallis test. F) Neither cytochalasin nor blebbistatin treatment affect osteoblast dedifferentiation measured as downregulation of *bglap*:GFP levels in segment 0 between 0 and 3 dpa. Plotted are the 3 dpa GFP levels relative to those in the same fish at 0 dpa. N (experiments) = 1, n (fins) = 10. Error bars represent 95% CI. Unpaired t-test. The observed relative difference is 4% (blebbistatin) and 16% (cytochalasin), the calculated smallest significant difference 38%, which is smaller than what we observe after retinoic acid treatment (Figure 3B, 63%). G) Inhibition of microtubule dynamics with nocodazole does not affect osteoblast migration. Images show overlay of 0 dpa (green) and 1 dpa (pink) pictures. N (experiments) = 1, n (fins) = 10, n (rays) = 20. Error bars represent 95% CI. Unpaired t-test. The observed relative difference is 1%, the calculated smallest significant difference 24% and thus smaller than what we observe in actomyosin treatment regimens (Figure 1D, 60 – 63%). Scale bar, 10 µm.

Besides the actomyosin network, microtubules (MT) are an important force to drive cell shape changes and cell motility (Etienne-Manneville, 2013). To analyse a potential role of MT in the amputation-induced migration of osteoblasts, we treated fish with nocodazole, a MT-binding drug that disrupts MT assembly/disassembly (Florian and Mitchison, 2016), using a regime which was shown to be effective in zebrafish (Poss *et al*., 2004). At 1 dpa, we could not detect a difference in the extent of migration in drug-treated fish compared to controls (Figure 2G). Together, these data indicate that actomyosin, but not microtubuli dynamics is required for injury-induced osteoblast migration.

### Osteoblast migration is independent of osteoblast dedifferentiation

We next wondered whether dedifferentiation is a prerequisite for osteoblasts to change their shape and to migrate. We have previously shown that NF-κB signalling negatively regulates osteoblast dedifferentiation (Mishra *et al*., 2020). To interfere with NF-κB signalling, we used genetic tools allowing for osteoblast-specific manipulation of the pathway. We induced expression of an inhibitor of NF-κB signalling (IkB super repressor, IkBSR) (Van Antwerp *et al*., 1996) in osteoblasts, using the tamoxifen-inducible Cre line *OlSp7*:CreERT2-p2a-mCherry^tud8^ (*osx*:CreER) (Knopf *et al*., 2011) and an ubiquitously expressed responder line *hsp70l*:loxP Luc2-myc Stop loxP nYPet-p2a-IκBSR*, cryaa*:AmCyan^ulm15Tg^, which expresses IkBSR and nYPet after recombination (*hs*:Luc to nYPet IκBSR). NF-kB signalling activation was induced by expressing constitutively active human IKK2 (IkB kinase) and nuclear localized BFP after recombination using *hsp70l*:loxP Luc-myc STOP loxP IKKca-t2a-nls-mTagBFP2-V5, cryaa:AmCyan^ulm12Tg^ (*hs*:Luc to IKKca BFP) fish (Mishra *et al*., 2020). Expression of GFP driven by the Cre-responder line *hsp70l*:loxP DsRed2 loxP nlsEGFP^tud9^ (*hs*:R to G) (Knopf *et al*., 2011) served as a negative control. Using these tools, we have previously shown that activation of NF-κB-signalling inhibits osteoblast dedifferentiation, while its suppression promotes their dedifferentiation (Mishra *et al*., 2020). Mosaic recombination allowed us to compare recombined and non-recombined cells within the same segment. Analysis of cell shape in the control line *hs*:R to G showed the elongation of both non-recombined and recombined cells in segment −1 compared to segment −2 (Figure 3A). Neither pathway inhibition by expression of IkBSR nor forced activation by expression of IKKca affected osteoblast elongation in segment −1. We conclude that NF-κB-signalling regulates osteoblast dedifferentiation, but not migration.

**Figure 3.**
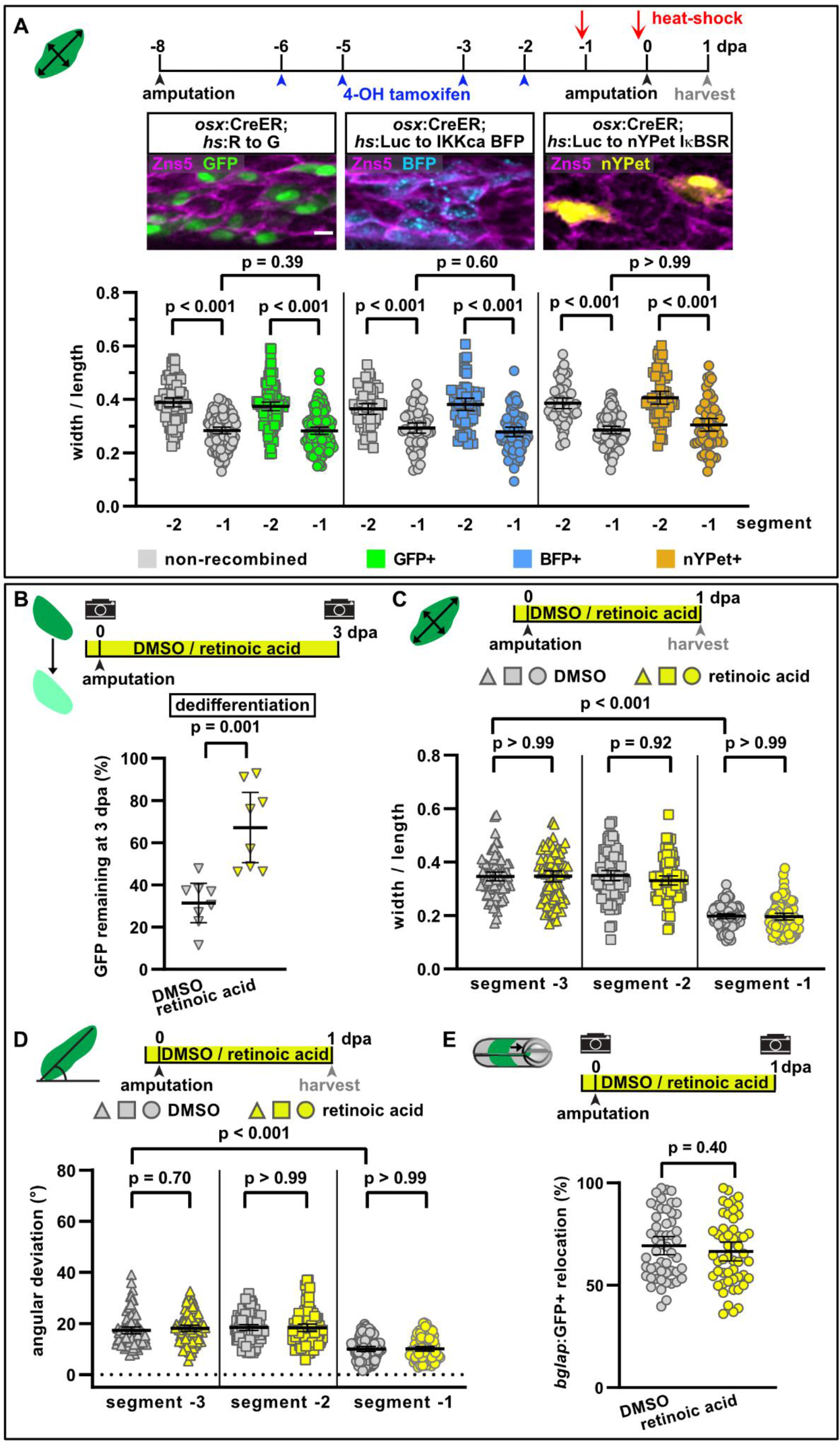
NF-kB and retinoic acid signalling regulate osteoblast dedifferentiation but not migration. A) Mosaic recombination in *osx*:CreER; *hs*:R to G, *osx*:CreER; *hs*:Luc to IKKca BFP and *osx*:CreER; *hs*:Luc to nYPet IκBSR fish. Zns5 labels the membrane of osteoblasts. Osteoblast roundness in recombined and non-recombined osteoblasts in segment −2 and −1 at 1 dpa. Recombined osteoblasts expressing IKKca (marked by BFP) or IκBSR (marked by nYPet) elongate in segment −1 to a similar extent as osteoblasts expressing the negative control GFP. N (experiments) = 1, R to G: n (fins) = 3, n (rays) = 10 n (cells) = 406; IKKca: n (fins) = 4, n (rays) = 7, n (cells) = 244; IκBSR: n (fins) = 8, n (rays) = 18, n (cells) = 246. Error bars represent 95% CI. Kruskal-Wallis test. Scale bar, 10 µm. B) Treatment with retinoic acid (RA) inhibits osteoblast dedifferentiation as measured by reduced downregulation of *bglap*:GFP levels in segment 0 between 0 and 3 dpa. N (experiments) = 1, n (fins) = 8. Error bars represent 95% CI. Unpaired t-test. C) Osteoblast roundness at 1 dpa. RA treatment does neither alter osteoblast cell shape in segments −3 and −2, nor elongation in segment −1. N (experiments) = 2, n (fins) = 10, n (rays) = 10, n (cells) = 83. Error bars represent 95% CI. Kruskal-Wallis test. The observed relative difference in segment −1 is 2%, the calculated smallest significant difference 7%, which is smaller than what we observe in cytochalasin treatment regimens (Figure 1A, 22%). D) Osteoblast orientation at 1 dpa. RA treatment does not affect osteoblast orientation in segments −3 and −2, nor alignment along the proximodistal axis in segment −1. N (experiments) = 2, n (fins) = 10, n (rays) = 10, n (cells) = 89. Error bars represent 95% CI. Kruskal-Wallis test. The observed relative difference in segment −1 is 1%, the calculated smallest significant difference 7%, which is smaller than what we observe in cytochalasin treatment regimens (Figure 1B, 32%). E) RA treatment does not affect bulk migration of osteoblasts toward the amputation plane in segment −1. N (experiments) = 3, n (fins) = 26, n (rays) = 52. Error bars represent 95% CI. Unpaired t-test. The observed relative difference is 2%, the calculated smallest significant difference 22%, which is smaller than what we observe in actomyosin treatment regimens (Figure 1D, 60 – 63%).

Retinoic acid signalling inhibits osteoblast dedifferentiation downstream of NF-κB-signalling (Blum and Begemann, 2015; Mishra *et al*., 2020). Treatment with retinoic acid (RA) impaired osteoblast dedifferentiation after amputation, as *bglap*:GFP downregulation in segment 0 was decreased (Figure 3B). In contrast, osteoblast elongation and alignment along the proximodistal axis in segment −1 at 1 dpa were not affected (Figure 3C, D). Similarly, RA did not affect bulk osteoblast migration towards the amputation plane (Figure 3E). Together, these data show that osteoblast dedifferentiation and migration are independently regulated, and they indicate that osteoblasts dedifferentiation is not a prerequisite for migration.

### The complement system is required for osteoblast migration in vivo

As shown above, we found that osteoblast cell shape changes and migration are directed towards the amputation site (Figure 1C). Directional cell migration is usually initiated in response to extracellular cues such as chemokines or signals from the extracellular matrix (Swaney, Huang and Devreotes, 2010). As part of the innate immune system, the complement system is activated immediately after injury, and activated complement factors have been shown to be able to act as chemoattractants for osteoblasts *in vitro* (Ignatius, Ehrnthaller, *et al*., 2011). We thus asked whether the complement system regulates osteoblast migration *in vivo*. The zebrafish genome contains homologs of all fundamental mammalian complement components (Boshra, Li and Sunyer, 2006; Zhang and Cui, 2014), including the central components C3 and C5 and the corresponding receptors C3aR and C5aR1, respectively. A second C5a receptor, C5aR2, has so far only been characterized in mammals, and sequence database interrogation indicates that it is absent in zebrafish. RNAscope in situ analysis revealed that *c5aR1* is expressed in mature osteoblasts (Figure 4A). Treatment with W54011, a specific C5aR1 antagonist (Sumichika *et al*., 2002), impaired osteoblast elongation in segment −1 at 1 dpa without affecting osteoblast cell shape in segments −2 and −3 (Figure 4B). As also the complement factor C3a can act as chemoattractant after trauma (Huber-Lang, Kovtun and Ignatius, 2013), we treated fish with SB290157, a specific antagonist of the complement receptor C3aR (Ames *et al*., 2001). Osteoblast elongation was impaired in segment −1 at 1 dpa, while cell shape in segment −2 and −3 was not affected (Figure 4C). Both drugs also significantly reduced bulk osteoblast migration (Figure 4D). To confirm these findings, we repeated the migration assay with PMX205, another C5aR1 antagonist (March *et al*., 2004; Jain, Woodruff and Stadnyk, 2013), which also reduced osteoblast migration at 1 dpa (Figure 4D). Together, these data strongly suggest that the complement system regulates injury-induced directed osteoblast migration. Interestingly, none of the complement inhibitors affected osteoblast dedifferentiation (Figure 4E). This supports the view that osteoblast dedifferentiation and migration are independent responses of osteoblasts to trauma. Complement components can also contribute to the regulation of cell proliferation and tissue regeneration (Mastellos and Lambris, 2002). Yet, regenerative growth was not affected in fish treated with either W54011, PMX205 or SB290157 (Figure 4 – figure supplement Figure 1A). Furthermore, inhibition of C5aR1 with PMX205 had no effect on osteoblast proliferation (Figure 4 – figure supplement 1B). We conclude that the complement system specifically regulates osteoblast migration, but not other responses of osteoblasts to trauma in zebrafish.

**Figure 4:**
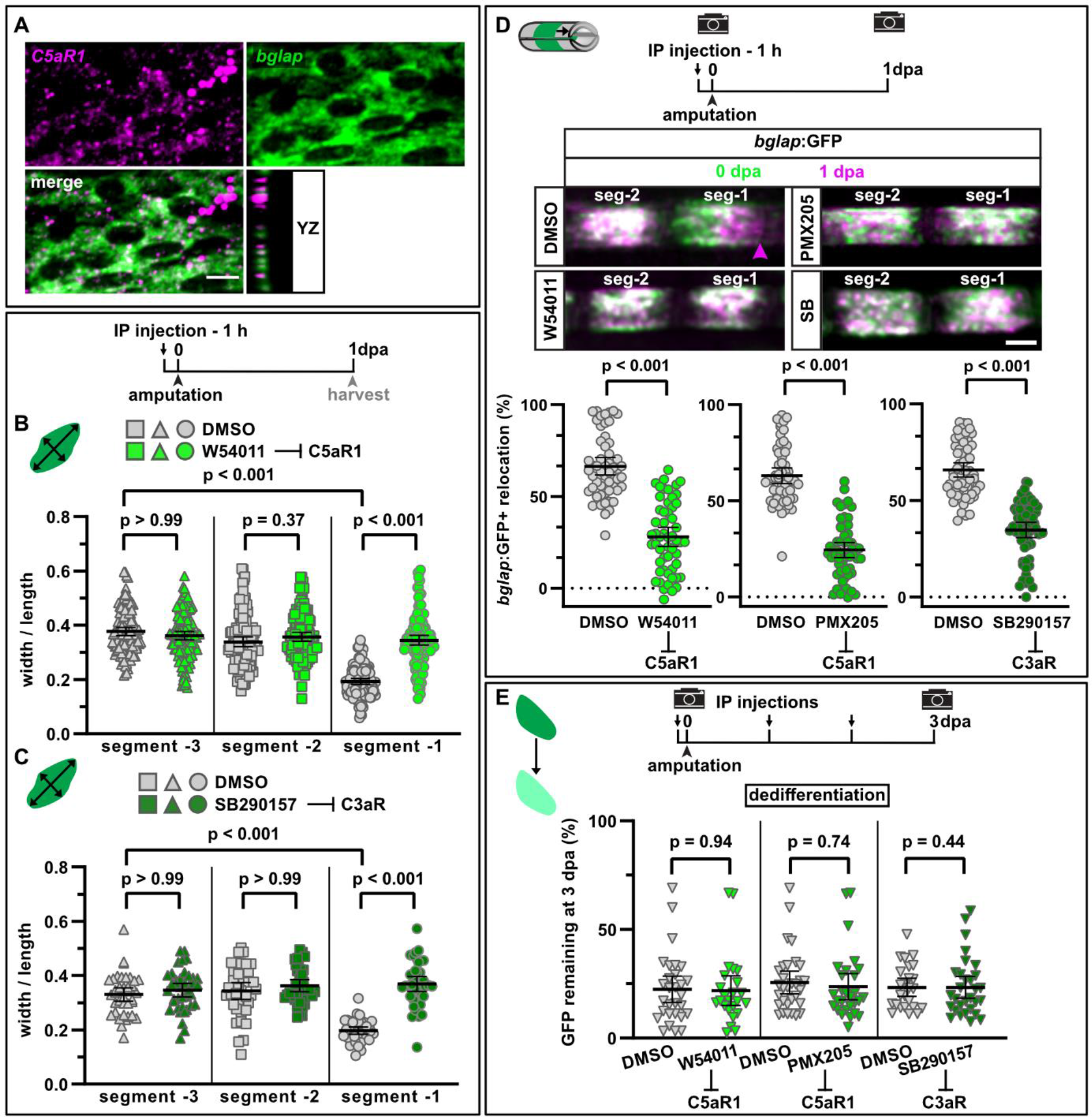
Complement system signalling is required for osteoblast cell shape change and migration after amputation. A) RNAscope in situ detection of *c5aR1* expression in *bglap* expressing osteoblasts in a non-injured segment. Scale bar, 10 µm. B) Osteoblast roundness at 1 dpa. The C5aR1 inhibitor W54011 does not alter osteoblast cell shape in segments −3 and −2, but cell elongation in segment −1 is inhibited. N (experiments) = 3, n (fins) = 5, n (rays) = 5, n (cells) = 116. Error bars represent 95% CI. Kruskal-Wallis test. C) Osteoblast roundness at 1 dpa. The C3R inhibitor SB290157 does not alter osteoblast cell shape in segments −3 and −2, but cell elongation in segment −1 is inhibited. N (experiments) = 1, n (fins) = 5, n (rays) = 5, n (cells) = 38. Error bars represent 95% CI. Kruskal-Wallis test. D) Inhibition of C5aR1 with either W54011 or PMX205 and inhibition of C3aR with SB290157 impairs bulk osteoblast migration. Images show overlay of 0 dpa (green) and 1 dpa (pink) pictures. N (experiments) = 3, W54011: n (fins) = 27, n (rays) = 54; PMX-205, SB290157: n (fins) = 29, n (rays) = 58. Error bars represent 95% CI. Mann-Whitney tests. Scale bar, 100 µm. E) Neither inhibition of C5aR1 (W54011, PMX205) nor of C3aR (SB290157) affects osteoblast dedifferentiation as measured by downregulation of *bglap*:GFP levels in segment 0 between 0 and 3 dpa. N (experiments) = 3, n (fins) = 31. Error bars represent 95% CI. Unpaired t-tests. The observed relative difference is 4% (W54011), 8% (PMX205) and 3% (SB290157), the calculated smallest significant differences are 35% (W54011), 31% (PMX205) and 31% (SB290157), which are smaller than what we observe after retinoic acid treatment (Figure 3B, 63 %).

Activation of the complement system is a general early response after injury that also occurs in wounds that heal, but do not trigger structural regeneration of the kind we observe after fin amputation. Thus, we wondered whether osteoblast migration can be triggered by injuries that do not induce structural regeneration. Lesions of the interray skin quickly heal (Chablais and Jaźwińska, 2010; Chen *et al*., 2015) but do not trigger bone regeneration. Yet, such wounds can induce signals that are sufficient to trigger structural regeneration in a missing-tissue context (Owlarn *et al*., 2017). We found that skin injuries induced close to the ray bone did not induce osteoblast migration, neither off the bone into the intraray, nor on the bone towards the joints (Figure 4 – figure supplement 1C). Osteoblasts also migrate towards fractures induced in the fin ray bone (Geurtzen *et al*., 2014). Interray injury combined with bone fracture resulted in osteoblast migration towards the fracture, but osteoblasts did not migrate away from the bone matrix into the interray skin (Figure 4 – figure supplement 2D). We conclude that generic wounding-induced signals are not sufficient to attract osteoblasts in the absence of additional pre-requisites, which might include the presence of a permissive substrate for migration.

### An internal bone defect model separates generic from regeneration-specific responses of osteoblasts to trauma

To separate potential generic injury responses from a regenerative response in bone, we established a bone injury model where we remove hemiray segments at two locations within one ray, leaving a centre segment with two injury sites, one facing proximally, the other distally (Figure 5A). It was recently shown that in a similar cavity injury model a blastema forms only at the distal-facing injury, while the proximally facing site displays no regenerative growth (Cao *et al*., 2021). In our hemiray removal model, both injury sites are equally severed from the stump, and thus cut off from innervation and blood supply. Therefore, any potential differences in the injury responses at the proximal vs distal injury site cannot be explained by differences in innervation or blood circulation. Within 2 days post injury (dpi), blood flow through the centre segment was restored (Figure 5 – figure supplement 1A, Video 2). At 3 dpi, *Runx2*-positive pre-osteoblasts and *Osterix-*positive committed osteoblasts accumulated almost exclusively in the defect beyond the distal injury, but not at the proximal injury (Figure 5B − D). Subsequently, only at the distal site, new bone matrix formed (Figure 5E). Thus, our hemiray removal injury model reveals that accumulation of a pre-osteoblast population and regeneration of bone occurs only at distal-facing wounds.

**Figure 5:**
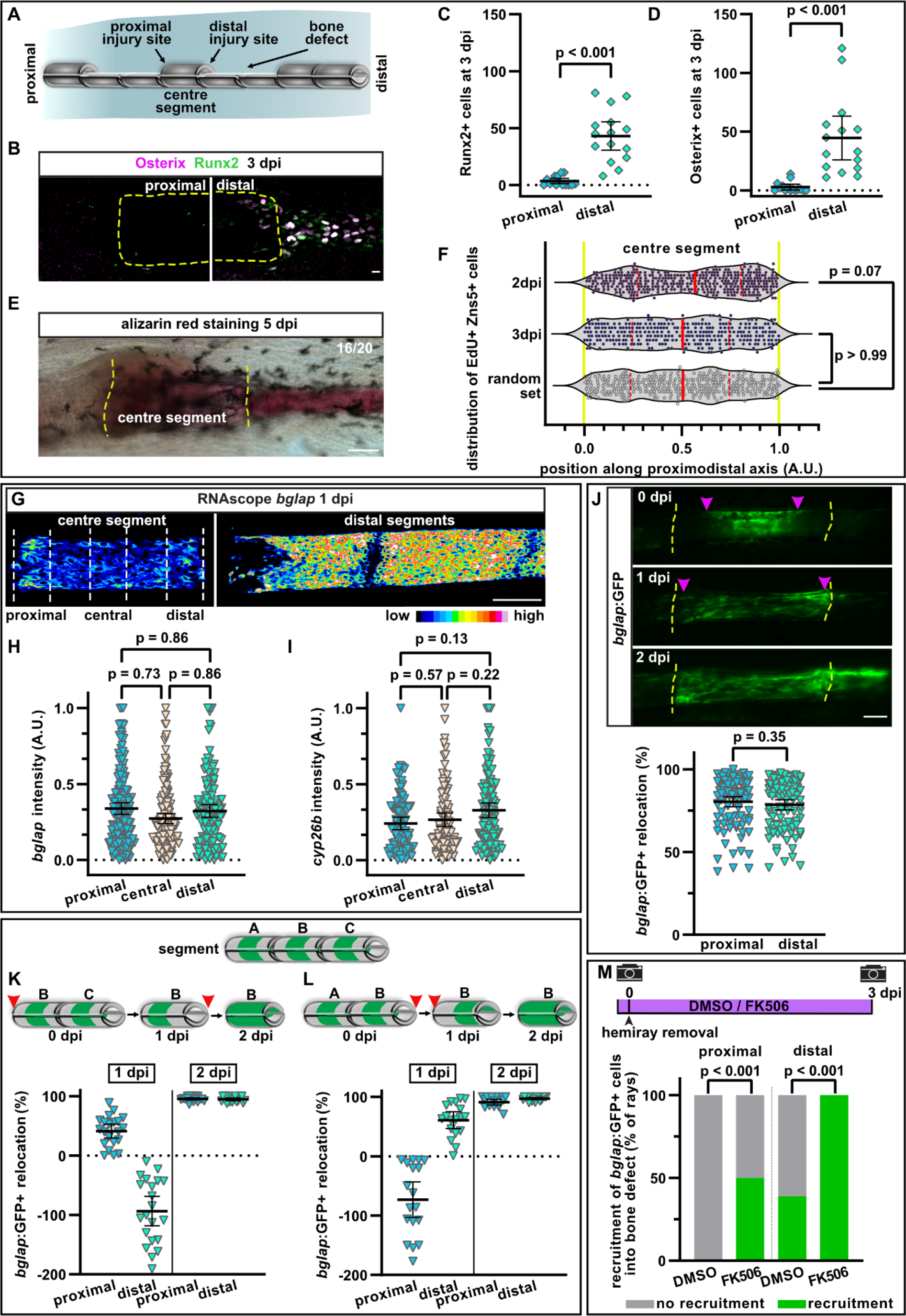
Hemiray removal results in polarized bone growth but symmetric osteoblast migration. A) Hemiray removal scheme. B) At 3 dpi, Runx2+ pre-osteoblasts and committed osteoblasts expressing Osterix have only accumulated in the bone defect beyond the distal injury site of the centre segment, but not beyond the proximal injury site. Dashed line outlines centre segment. Scale bar, 10 µm. C) Quantification of the number of Runx2+ cells located in the bone defect beyond the proximal and distal injury sites of the centre segment at 3 dpi. n (rays) = 15. Error bars represent 95% CI. Wilcoxon matched-pairs test. D) Quantification of the number of Osterix+ cells located in the bone defect beyond the proximal and distal injury sites of the centre segment at 3 dpi. n (rays) = 15. Error bars represent 95% CI. Wilcoxon matched-pairs test. E) At 5 dpi, mineralized bone as detected by alizarin red staining has formed beyond the distal, but not the proximal injury site of the central segment. Dashed line indicates centre segment. n = 16/20 rays with distal bone formation. Scale bar, 100 µm. F) Distribution of all positions along the proximodistal axis of the centre segment where proliferating EdU+ Zns5+ osteoblasts have been observed at 2 and 3 dpi. Yellow dashed lines indicate segment border. Solid red lines, median; dashed lines, quartiles. 2 dpi: n (rays) = 13, n (cells) = 306; 3 dpi: n (rays) = 17, n (cells) = 305; random set: n (groups) = 12, n (points) = 360. Kolmogorov-Smirnov test. G) RNAscope in situ detection of *bglap* expression in the centre segment and the two adjacent segments distal to the bone defect at 1 dpi. Dashed lines indicate the proximal, central and distal regions of the centre segment used for quantification. Scale bar, 100 µm. H) Single cell analysis of *bglap* RNAscope intensity relative to the brightest signal in proximal, central and distal regions of the centre segment at 1 dpi. n (segments) = 8. Error bars represent 95% CI. Holm-Sidak’s multiple comparison test. I) Single cell analysis of *cyp26b1* RNAScope signal intensity relative to the brightest signal in proximal, central and distal regions of the centre segment at 1 dpi. n (segments) = 7. Error bars represent 95% CI. Holm-Sidak’s multiple comparison test. J) Migration of *bglap*:GFP+ osteoblasts towards both injury sites of the centre segment. Yellow dashed lines, segment borders. Pink arrowheads indicate relocation of GFP+ osteoblasts. Distal to the right. n (segments) = 95. Error bars represent 95% CI. Wilcoxon matched-pairs test. Scale bar, 100 µm K, L) Sequential hemiray injuries reveal no preference for osteoblast migration in distal or proximal directions. Relocation of *bglap*:GFP+ osteoblasts in the centre segment (segment B). Red arrowheads indicate timepoints at which the adjacent hemirays (A = proximal, C = distal) were removed. Negative relocation indicates increased distance between GFP+ cells and the joint. K) Relocation of *bglap*:GFP+ osteoblasts in the centre segment (segment B). Removal of the proximal adjacent segment A at 0 dpi, followed by removal of the distal adjacent segment C at 1 dpi. L) Relocation of *bglap*:GFP+ osteoblasts in the centre segment (segment B). Removal of the distal adjacent segment C at 0 dpi, followed by removal of the proximal adjacent segment A at 1 dpi. M) Migration of osteoblasts into the bone defect at 3 dpi. Treatment with FK506 induces recruitment of GFP+ cells beyond the proximal injury site and increases the number of centre segments that show recruitment of GFP+ cells beyond the distal injury site. N (experiments) = 1, n (segments) = 56 (DMSO), 17 (FK506). Error bars represent 95% CI. Fisher’s test.

In response to regular fin amputations, the initiation of osteoblast dedifferentiation, migration and proliferation occur prior to the accumulation of a pre-osteoblast population in the blastema. Thus, we asked whether any of these osteoblast injury responses occurs at proximal-facing injuries in the hemiray injury model, which fail to form a blastema containing pre-osteoblasts. At 2 and 3 dpi, proliferation of osteoblasts was observed throughout the centre segment (Figure 5F). While at 2 dpi the median of the distribution of proliferating osteoblasts along the proximodistal axis was slightly shifted distally compared to a random distribution set, at 3 dpi proliferation was equally distributed along the segment (Figure 5F). Thus, the polarized regenerative response is not reflected by an equally polarized osteoblast proliferation.

We next wondered whether the polarized bone regeneration is due to varying degrees of osteoblast dedifferentiation along the segment. We analysed the expression of *bglap* using RNAScope in situ hybridization on single cell level in a proximal, central and distal region of the centre segment at 1 dpi. Within such a distance (∼ 300 µm), a gradual change in *bglap* intensity can be observed after amputation (Figure 1 – figure supplement 1F). Yet, we could not detect differences in the extent of *bglap* expression between the regions of the centre segment (Figure 5G, H), indicating that both injury sites induced osteoblast dedifferentiation. We also analysed the expression of *cyp26b1*, an enzyme involved in retinoic acid signalling, which is upregulated in dedifferentiating osteoblasts (Blum and Begemann, 2015). *Cyp26b1* was upregulated along the entire segment, with no enrichment at the distal injury (Figure 5I). Together, these data show that osteoblast dedifferentiation occurs to a similar extent in the bone facing the proximal and distal injury.

Intriguingly, osteoblasts in the centre segment also migrated towards both amputation planes, ultimately spreading across the entire segment (Figure 5J). The magnitude of migration was similar towards both wound sites. This suggests that at this early timepoint, there is no hierarchy between the injury sites, and that osteoblasts react to an initial wound signal, which is independent of the proximodistal position of the injury. Furthermore, as osteoblasts also migrate towards the proximal site where no blastema will form, it appears that osteoblasts migrate independently of whether this will be followed by bone regeneration. To further test this, we temporally separated the removal of the two hemirays at the two sides of the centre segment (Figure 5K, L). We first extracted only one hemiray (segment A) and analysed osteoblasts of the adjacent distal segment (segment B). Within one day, the distance between osteoblasts and the distal segment border increased, while proximally it decreased, indicating a directed migration of the osteoblasts towards the proximally facing injury (Figure 5K). We then removed the hemiray of segment C (the one located distally to the centre segment B), and analysed osteoblast locations in segment B one day later. The distance between osteoblasts and the proximal segment border further decreased, indicating that the osteoblasts continued to migrate towards the proximal injury (Figure 5K). However, at the distal side, the previously increased distance was decreased, suggesting that osteoblasts reversed their direction and migrated towards the distal injury as well (Figure 5K). This phenomenon is independent of the order of hemiray removal: when the distal adjacent hemiray (segment C) was removed first, osteoblast migrated distally (Figure 5L). When we then removed the proximal adjacent hemiray (segment A), the distal osteoblasts continued to migrate distally, while proximally located osteoblast migrated towards the new injury site, even though at this site no blastema will form (Figure 5L). These data strongly suggest that all bone injuries release signals that attract osteoblasts and that osteoblasts are equally likely to migrate proximally and distally. Thus, the lack of regenerative growth and bone formation at proximally-facing injuries cannot be explained by an intrinsic polarity or bias in the migration of osteoblasts on the bone matrix along the proximodistal axis of the fin.

At 2 dpi, at the distal-facing injury site, *bglap*:GFP+ osteoblasts of the centre segment accumulated beyond the bone matrix in the defect, where a blastema appears to form (Figure 5J). However, no osteoblasts outside the bone matrix could be observed at the proximal side of the centre segment at 3 dpi (Figure 5J). Thus, while osteoblasts migrate along the bone matrix towards both injury sites, they exclusively migrate beyond the matrix at the distal-facing injury. In a cavity injury model, Cao et al. have shown that inhibition of the Ca^2+^/calmodulin dependent phosphatase calcineurin can induce blastema formation at proximal-facing injuries (Cao *et al*., 2021). To analyse if such an intervention can also trigger osteoblast migration beyond the bone matrix, we treated fish with the calcineurin inhibitor FK506. Indeed, we could observe recruitment of *bglap*:GFP+ cells into the defect at the proximal amputation site in FK506 treated fish (Figure 5M). Noteworthy, also migration beyond the distal amputation site was enhanced. Therefore, to test if calcineurin inhibition specifically enabled migration of osteoblasts outside of the bone matrix or generally facilitated migration, we analysed osteoblast migration after regular fin amputation. FK506 treatment enhanced osteoblast migration at 1 dpa (Figure 5 – figure supplement 1B), indicating that calcineurin might generally negatively regulate osteoblast migration, and does not specifically facilitate distally directed regenerative responses.

The lack of osteoblast accumulation at proximally-facing injuries could be due to absence of chemical or mechanical cues that allow them to migrate into the defect. Alternatively, migration could be actively inhibited or osteoblasts could be eliminated at proximal injuries. One possibility would be increased osteoblast apoptosis at proximally-facing injuries. However, we could not detect any apoptotic osteoblasts at all in the first two days after injury (data not shown), and also at 3 dpi the number of apoptotic osteoblasts was negligible at both the proximal and distal injury sites (Figure 5 – figure supplement 1C). Another possibility would be that osteoclasts interfere with osteoblast differentiation and bone formation at the proximal injury. To analyse if more osteoclasts are recruited to the proximal injury in the hemiray removal model, we analysed the distribution of *cathepsinK*:YFP+ cells, a marker for osteoclasts, after hemiray removal. However, osteoclasts accumulated at both wound sites by 3 dpi (Figure 5 – figure supplement 1D).

Bone is thicker at the proximal base of the fin than at its distal end (Marí-Beffa and Murciano, 2010; Pfefferli and Jazwinska, 2015). We thus wondered whether this structural heterogeneity along the fin ray could determine the different regenerative outcomes at proximal vs distal facing injuries. Yet, we did not observe a measurable difference in the diameter, radius and thickness of one hemiray segment at its proximal vs distal end (Figure 5 – figure supplement 1E). This suggests that structural heterogeneities of the bone along the proximodistal axis are too small at the scale of the centre segment to explain the radically different regenerative outcome at the proximal vs distal injury in the hemiray removal model.

## Discussion

### Generic and regeneration-specific response of osteoblasts to trauma

In this study, we interrogate the interrelatedness of several *in vivo* responses of osteoblasts to trauma using the zebrafish fin model. After fin amputation, osteoblasts change their cell shape – they elongate and orient along the proximodistal axis of the fin – and migrate distally towards the amputation plane (Figure 6A). In addition, osteoblasts dedifferentiate, that is they downregulate expression of the differentiation marker *bglap* and initiate expression of the pre-osteoblast marker *runx2*. Finally, they proliferate and migrate off the bone surface to found a population of osteogenic progenitors within the regeneration blastema that drives bone regeneration. Surprisingly, we find that these trauma responses appear to occur largely independently of each other. In particular, osteoblast dedifferentiation does not seem to be a requirement for osteoblast cell shape change and migration. These conclusions are based on two lines of evidence: first, we identified molecular interventions that can interfere with one process without affecting the others: Retinoic acid and NF-κB signalling pathways regulate only osteoblast dedifferentiation, but not cell shape changes and migration, while the complement and actomyosin systems are required for osteoblast cell shape change and migration, but not for dedifferentiation (Figure 6A). Secondly, using a hemiray injury model, where a proximally and distally facing bone injury is introduced in the same fin ray, we find that osteoblast migration towards the injury occurs at both sites (Figure 6B). Similarly, osteoblasts dedifferentiate equally along the centre segment. Stunningly however, osteoblast migration beyond the bone into the injury defect, formation of a blastema containing a pre-osteoblast population, and regenerative bone growth only occurs at the distal-facing injury. Thus, osteoblast migration and dedifferentiation represent generic injury responses that can occur independently of a regenerative response (Figure 6B).

**Figure 6.**
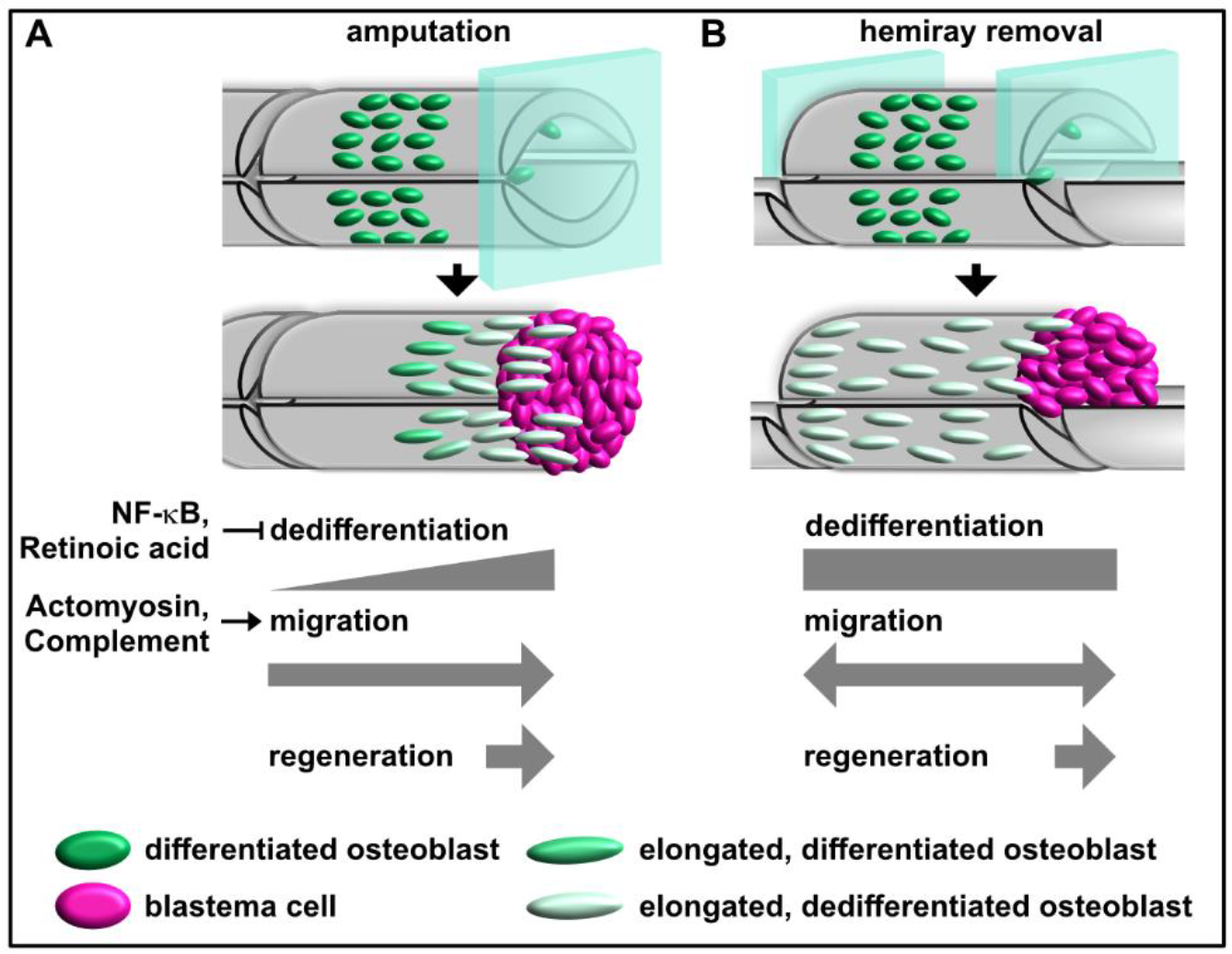
Model for osteoblast responses to fin amputation (A) and hemiray removal (B). Osteoblast dedifferentiation and migration represent generic injury responses that are differentially regulated and can occur independently of each other and of regenerative bone growth. A) After amputation, osteoblasts downregulate the expression of differentiation markers. The extent depends on their distance to the amputation site, with osteoblasts close to the amputation site displaying more pronounced dedifferentiation. In addition, osteoblasts elongate and migrate towards the amputation plane and beyond into the blastema, which forms at the amputation site. Thus, osteoblast dedifferentiation, migration and bone regeneration are all oriented distally. Osteoblast dedifferentiation is negatively regulated by NF-κB and retinoic acid signalling, while actomyosin dynamics and the complement system are required for directed osteoblast migration. B) In the hemiray removal model, a proximal and a distal injury are created on both sides of a remaining centre segment. The extent of osteoblast dedifferentiation is even along the centre segment, and osteoblasts migrate towards both injury sites. Yet, only at the distal-facing site, osteoblasts migrate into the bone defect, and blastema formation and bone regeneration only occur here.

The fact that dedifferentiation and migration do not always result in bone regeneration raises questions about their function. We have previously shown that experimentally enhanced dedifferentiation results in an increased population of progenitor cells in the regenerate, which however had detrimental effects on bone maturation (Mishra *et al*., 2020), indicating that dedifferentiation must be kept within certain limits. Unfortunately, currently existing tools to block dedifferentiation are either mosaic (activation of NF-κB signalling using the Cre-lox system) or cannot be targeted to osteoblasts alone (treatment with retinoic acid). Thus, we can currently not yet test what consequences specific, unmitigated perturbation of osteoblast dedifferentiation has for overall fin / bone regeneration. Conversely, the interventions presented here that specifically perturb osteoblast migration act only transiently, that is they can severely delay, but not fully block migration. Thus, an unequivocal test of the importance of osteoblast migration for bone regeneration likewise requires different tools. However, it has been shown that fin bone can regenerate even after genetic ablation of osteoblasts, due to activation of non-osteoblastic cells which can drive *de novo* osteoblast differentiation (Singh, Holdway and Poss, 2012; Ando *et al*., 2017). We speculate that the robustness of bone regeneration in zebrafish is enhanced by redundant mechanisms of source cell formation, which can either derive from pre-existing osteoblasts, which dedifferentiate and migrate towards the injury, or from (reserve) stem / progenitor cells. Dedifferentiation and migration in response to injuries that eventually do not trigger bone regeneration likely adds to the robustness, since it primes all injuries for regeneration. Interestingly, work in planaria and zebrafish fins has previously shown that all wounds, irrespective of whether they trigger (structural) regeneration or result merely in wound healing, appear to activate a generic signalling response (Wurtzel *et al*., 2015; Owlarn *et al*., 2017). The differential outcome (structural regeneration vs wound healing) is rather determined further downstream, and might dependent on systems that measure how much anatomy is missing (Owlarn *et al*., 2017). In the present work we show that this principle also applies to osteoblast trauma responses. It appears that generic wounding-induced signals trigger the generic responses of osteoblast dedifferentiation and migration, while the decision of whether these will be followed by bone regeneration is controlled by other determinants further downstream. Currently, the molecular nature of such determinants that can sense the amount of missing anatomy and the difference between proximally and distally-facing injuries remains rather mysterious.

### Osteoblast migration requires actomyosin, but not microtubule dynamics

The ability of cells to migrate is essential for many processes such as embryonic development, immune surveillance and wound healing. Cell motility is a highly dynamic process, and the cytoskeleton is an essential component in creating motility-driving forces and coordinated movement. Before migration, cells often display a roundish morphology without distinct protrusion, while the migrating cell extends protrusions oriented along the axis of migration (Aman and Piotrowski, 2010). We observe such a change in cell shape in osteoblasts close to the amputation plane, which become directionally elongated after amputation. Forces generated by the actin cytoskeleton, both through its ability to rapidly assemble and disassemble, and actomyosin-driven contraction, are essential regulators of cell shape change and migration. Additionally, microtubules (MT) can modulate migration by contributing to the formation of cell protrusions, and through their role in signalling and cellular trafficking (Etienne-Manneville, 2013). However, the involvement of MT is cell-type dependent, as migration of keratocytes and neural cells is independent of MTs (Etienne-Manneville, 2013). For osteoblast migration, the underlying cytoskeletal dynamics are still poorly understood. The reorganization of actin and MT is mediated by various regulators such as binding proteins and Rho GTPases, and interfering with their function provide some evidence on the involvement of the respective cytoskeletal element in osteoblasts. siRNA-mediated knockdown of the actin-monomer binding protein Profilin in osteoblastic MC3T3 cells reduces their motility in *in vitro* migration assays (Miyajima *et al*., 2012). Rho-associated protein kinase ROCK is a downstream effector of the small GTP-binding protein Rho and mediates, among others, cell contraction and migration. ROCK inhibition induces rearrangement of the actin cytoskeleton and stimulates migration in cultured human osteoblasts (Zhang *et al*., 2011). While these data indicate the importance of actomyosin dynamics, also MT might be involved. MACF1 (microtubule actin crosslinking factor 1), which can bind to both actin and MT, promotes MT stability and dynamics. MACF1 overexpression in MC3T3-E1 cell enhances migration *in vitro* and *in vivo*, while migration is reduced in sh-MACF1 cells (Su *et al*., 2020, 2022). Our data imply that for osteoblasts in the zebrafish fin, cell elongation and motility strongly depend on the dynamic actomyosin network, while microtubules are dispensable.

### The complement system regulates osteoblast migration in vivo

Directed osteoblast migration is an important step in bone remodelling during bone tissue homeostasis and during bone fracture healing (Thiel *et al*., 2018). Cell migration can be directed by gradients of ligands acting as chemoattractants. For osteoblasts, over 20 different factors which can elicit a migratory response in *in vitro* assays have been identified (Dirckx, Van Hul and Maes, 2013; Thiel *et al*., 2018). However, confirmation of their roles in *in vivo* models is still sparse, largely due to the difficulty in assaying osteoblast migration independently of other phenotypes, e.g. osteoblast proliferation or differentiation, in rodents *in vivo*. Indirect evidence is given for PDGFRβ, where callus formation after fracture is reduced in mice lacking PDGFRβ in osteoprogenitors, suggesting that the mobilization of osteoprogenitors requires PDGFRβ signalling (Böhm *et al*., 2019). In TGF-β knockout mice, injected bone marrow stem cells (BMSCs) fail to home to trabecular bone surfaces, as they do in wild-type mice (Tang *et al*., 2009). Similarly, VEGF and BMP-2 induced homing of BMSCs in mice and rabbits (Zhang *et al*., 2014). Yet, direct observation of migrating osteoblasts is lacking in these studies. Using live zebrafish and time lapse imaging, we show that the complement system regulates osteoblast migration *in vivo*. The complement cascade is a classical column of innate immunity. The liver is the main producer of complement proteins, and the soluble effector proteins circulate in serum and interstitial fluid as precursors (Merle *et al*., 2015). Activation of complement can occur via different pathways, all of which result in a sequential cascade of enzymatic reactions eventually leading to the cleavage of the complement proteins C3 and C5 into the potent anaphylatoxins C5a and C3a. As part of their function during the inflammatory response, C3a and C5a act as chemoattractants which can recruit immune cells such as mast cells, monocytes and granulocytes (Arbore, Kemper and Kolev, 2017). The complement receptors C3aR and C5aR are expressed in mammalian osteoblasts (Ignatius, Schoengraf, *et al*., 2011), and C3a and C5a act as chemokines for osteoblasts *in vitro* (Ignatius, Ehrnthaller, *et al*., 2011). Therefore, the complement system might act as modulator of osteoblast migration during bone remodelling and regeneration. Here we provide support for this hypothesis *in vivo*. Interestingly, our data indicate that both C3a and C5a act as guidance cues for osteoblasts, ensuring the recruitment of osteoblasts to the injury plane.

There is growing evidence that complement factors can modulate highly diverse processes such as cell growth, differentiation and regeneration in various tissues. Concurrently, components of the complement system display distinct expression profiles in different tissues and can be found in such divergent cell types such as adipocytes, astrocytes, fibroblasts and endothelial cells (Mastellos and Lambris, 2002; Arbore, Kemper and Kolev, 2017). Both C3 and C5 were shown to play a role during liver regeneration in mammals. C3^-/-^ as well as C3R^-/-^ deficient mice show impaired liver regeneration (Markiewski *et al*., 2004). C5^-/-^ deficient mice suffer from defective liver regeneration, with hepatocytes displaying diminished mitotic activity (Mastellos *et al*., 2001). As inhibition of C5aR1 impairs hepatocyte proliferation (Mastellos *et al*., 2001), and expression of C5aR1 is upregulated during liver regeneration (Daveau *et al*., 2004), C5 might act by regulating hepatocyte proliferation after liver injury. Regulation of cellular proliferation by complement proteins has been demonstrated in a broad range of tissues (Mastellos and Lambris, 2002). Expression of *c5aR1* is upregulated in the regenerating hearts of zebrafish, axolotls, and neonatal mice, and receptor inhibition impairs cardiomyocyte proliferation (Natarajan *et al*., 2018). Noteworthy, in our study on zebrafish fin regeneration, inhibition of C5aR1 had no effect on osteoblast proliferation and regenerative growth. The distinct outcome of complement activation might be depending on the location of complement factors. Intracellular C3 and C5 storage has been proposed in T cells, and the respective cleavage products C3a and C5a bind to intracellular receptors (Liszewski *et al*., 2013; Arbore *et al*., 2016). In human CD4+ T cells, intracellular C3a regulates cell survival, while activation of membrane-bound C3aR modulates induction of the immune response (Liszewski *et al*., 2013). In addition to the receptors, both C3 and C5 are expressed in mammalian osteoblasts (Ignatius, Schoengraf, *et al*., 2011). Yet, as we observe directed migration of osteoblasts towards the injury site after zebrafish fin amputation, it is likely that a gradient of C3a and C5a, caused by local expression or activation of C3 and C5, respectively, at the injury site, is dominant over any complement ligands that are produced in osteoblasts themselves. Intriguingly, during newt and urodele limb regeneration, C3 is expressed in blastema cells, which implies a role in mediating regeneration (Del Rio-Tsonis *et al*., 1998; Kimura *et al*., 2003). However, as during zebrafish fin regeneration osteoblasts migrate before blastema formation, a different cellular source likely produces C3 and C5. Yet, we cannot exclude that complement emanating from the blastema is responsible for the migration of osteoblasts beyond the bone context at later stages, which we only observed at sites where a blastema forms. Prior to blastema formation, the injury site is covered by a wound epidermis. During newt limb regeneration, C5 was shown to be absent from the blastema but highly expressed in the wound epidermis (Kimura *et al*., 2003). Thus, for the early migration of osteoblasts to the injury, complement activation in epidermal cells might be accountable. Such an injury-induced response would be locally activated at both sides in the hemiray removal model, corresponding with the symmetric migration we observed. Osteoblast migration might additionally be modulated by surface recognition features. In the dermal bones of the fin, osteoblasts line the bone matrix as a single layer, suggesting that osteoblasts might recognize matrix components such as collagens. Osteoblasts might utilize surface features as landmarks, such that the osseous surface serves as navigation signal for osteoblast migration, while additional signals are necessary for the migration out of the bone context. Here, specific signals emanating from the blastema might be necessary, as we also do not observe osteoblast migration towards interray injuries, where no blastema forms. Further analysis identifying the source of complement factors during zebrafish fin regeneration will help to understand the regulation of directed osteoblast migration.

### Determinants of regeneration-specific osteoblast trauma responses

Removing part of the bone within a ray results in polarized bone regeneration that is exclusively directed towards the distal end of the fin. This polarized regenerative response is already evident within 2 days post injury. In a recently published zebrafish fin cavity injury model, blastema markers were shown to be solely upregulated at the distal facing injury (Cao *et al*., 2021). In our hemiray-removal model, we analyse the proximal and distal injury of a centre segment. Thus, both injury sites are equally severed from innervation and blood supply, giving us confidence that the differential regenerative response at proximal and distal injuries is not due to differences in circulation or innervation. We show that osteoblasts migrate from the bone matrix into the injury only at the distal-facing injury. Consequently, a population of Runx2+ and Osterix+ (pre-) osteoblasts only accumulates beyond the distal injury. The distally-oriented polarized regenerative response implies intrinsic mechanisms regulating diverse outcome from seemingly equal injuries (the removal of the adjacent hemiray). One previously suggested regulator of distally-oriented regeneration is the Ser/Thr phosphatase calcineurin (Cao *et al*., 2021). A positive role of calcineurin on bone formation was shown previously *in vitro* and *in vivo* in mice (Sun *et al*., 2005), attributing its effect to increased osteoblast differentiation. During zebrafish fin regeneration, inhibition of calcineurin increases regenerative growth (Kujawski *et al*., 2014). Importantly, the polarized, distally-oriented blastema formation in the cavity injury model can be overridden by calcineurin inhibition, which induces blastema formation also at the proximal injury (Cao *et al*., 2021). We also found that calcineurin inhibition can trigger the recruitment of osteoblasts into the bone defect at the proximal injury site in our hemiray removal model. Yet, we also observed increased distally-oriented osteoblast migration upon FK506 treatment within the bone segment after amputation, suggesting that calcineurin does not specifically regulate the polarity of trauma responses along the proximodistal axis, but generally dampens trauma responses at all injuries, including osteoblast migration and cell proliferation. The role of calcineurin in cell migration is still controversial. Overexpression of the negative calcineurin regulator RCAN1 reduces migration of various cancer cell lines in vitro (Espinosa *et al*., 2009), while overexpression of calcineurin Aα increases migration of small cell lung cancer cells *in vitro* (Liu *et al*., 2010). In contrast, inhibiting calcineurin activity by FK506 increases migration in melanocytes and melanoma cells *in vitro* (Jung and Oh, 2016).

Our results imply that the same population of osteoblasts responds to bone injury by dedifferentiation and migration. Interestingly, *in vitro* treatment of osteocytes with parathyroid hormone (PTH) results in increased migratory responsiveness, while the cells lose their mature phenotype (Prideaux *et al*., 2015), which might be an indication for their dedifferentiation (Torreggiani *et al*., 2013; Thiel *et al*., 2018). *In vitro*, migration of undifferentiated osteoblast progenitor cells is more efficient than migration of mature, differentiated osteoblasts (Thiel *et al*., 2018). Yet, surprisingly, our data suggests that in the zebrafish fin, migration of osteoblasts is independent from their differentiation state, as inhibition of dedifferentiation did not affect osteoblast migration. Neither C3a nor C5a affect osteogenic differentiation of human mesenchymal stem cells (MSC) (Ignatius, Schoengraf, *et al*., 2011). Similarly, we found that the complement system does not regulate osteoblast dedifferentiation in the zebrafish. Our findings that osteoblast dedifferentiation and migration are regulated by different molecular signals and thus can be uncoupled from each other show that dedifferentiation is not a prerequisite for osteoblast migration.

In conclusion, our findings support a model in which bone regeneration involves both generic and regeneration-specific trauma responses of osteoblasts. Morphology changes and directed migration towards the injury site as well as dedifferentiation represent generic responses that occur at all injuries even if they are not followed by regenerative bone formation. While migration and dedifferentiation can be uncoupled and are (at least partially) independently regulated, they appear to be triggered by signals that emanate from all bone injuries. In contrast, migration off the bone matrix into the bone defect, formation of a population of (pre-) osteoblasts and regenerative bone formation represent regeneration-specific responses that require additional signals that are only present at distal-facing injuries. The identification of molecular determinants of the generic vs regenerative responses will be an interesting avenue for future research.

## Methods and materials

### Animals

All procedures involving animals adhered to EU directive 2010/63/EU on the protection of animals used for scientific purposes, and were approved by the state of Baden-Württemberg (Project numbers 1193 and 1494) and by local animal experiment committees. Fish of both sexes were used. Housing and husbandry followed the recommendations of the Federation of European Laboratory Animal Science Associations (FELASA) and the European Society for Fish Models in Biology and Medicine (EUFishBioMed) (Aleström *et al*., 2020).

The following pre-existing transgenic lines were used: *bglap*:GFP (*Ola.Bglap.1*:EGFP^hu4008^) (Vanoevelen *et al*., 2011), *osx*:CreER (*Ola.Sp7*:CreERT2-p2a-mCherry^tud8^) (Knopf *et al*., 2011), *entpd5*:kaede (TgBAC(entpd5a:Kaede)) (Huitema *et al*., 2012), cathepsinK:YFP (Apschner *et al*., 2014), *hs*:R to G (*hsp70l*:loxP DsRed2 loxP nlsEGFP^tud9^) (Knopf *et al*., 2011), *hs*:Luc to IKKca BFP (*hsp70l*:loxP Luc-myc STOP loxP IKKca-t2a-nls-mTagBFP2-V5, *cryaa*:AmCyan^ulm12Tg^) (Mishra *et al*., 2020). To create the line *hsp70l*:loxP Luc2-myc STOP loxP nYPet-p2a-IκBSR*, cryaa*:AmCyan^ulm15Tg^, in short *hs*:Luc to nYPet IκBSR, the following elements were assembled by Gibson assembly and restriction-based cloning methods: MiniTol2 inverted repeat, attP site, zebrafish *hsp70l* promoter, loxP, firefly luciferase 2, 6x myc tag, ocean pout antifreeze protein polyA signal, loxP, nYPet, p2a, IκBSR (IkappaB super-repressor), SV40 polyA signal, zebrafish *cryaa* promoter, AmCyan, SV40 polyA signal, MiniTol2 inverted repeat. IkBSR is a mutated version of mouse IKappa B alpha (Entrez gene, Nfkbia A1462015), described in (Van Antwerp *et al*., 1996), whose inhibitory N-terminal Serine residues 32 & 36 are mutated to Alanine. Phosphorylation sites in the C-terminal PEST domain are also mutated: Serines 283, 288, 293 are converted to Alanine; Threonines 291, 296 also to Alanine and Tyrosine 302 to Aspartic acid. Tol2 mediated transgene insertion was used to create a stable transgenic line. One subline was selected based on widespread expression after heat-shock in the adult fin and efficient recombination in embryos when crossed with an ubiquitin-promoter driven Cre driver line (unpublished).

To achieve optimal expression levels of the *osx*:CreER transgene, we activated its expression in adult fins as described previously (Mishra *et al*., 2020). Briefly, fish were amputated and allowed to regenerate for 8 days, at which time-point mineralized bone has formed in the regenerate. We then re-amputated through the mineralized part of the regenerate and analysed the stump of this second amputation.

### Fin amputations and hemiray removal

Adult zebrafish were anaesthetized with 625 µM tricaine, and the caudal fins were amputated through the 2^nd^ segment proximal to the bifurcation. For hemiray removal, small surgical blades were used to cut through four joints proximal to the bifurcation in the 2^nd^ ray from ventral and dorsal, and fine tweezers were used to remove the upper, left hemiray (facing towards the experimenter) of the segments located proximal and distal of one central segment. For analysis of migration and dedifferentiation, one hemiray segment was removed on either side. For analysis of proliferation and Runx2 and Osterix expression, two hemiray segments were removed on both sides. For alizarin red staining at 5 dpi, three hemiray segments were removed. Fish were allowed to regenerate at 27-28.5°C.

### Pharmacological interventions

10 µl of the following drugs (in PBS) were injected intraperitoneally (IP) using a 100 µl Nanofil syringe (World Precision Instruments NANOFIL-100): blebbistatin 750 nM, cytochalasin D 30 µM, W54011 10 µM, SB290157 10 µM, PMX205 10 µM, R115866 0.67 mM. Fish were injected once daily, starting 1h prior to injury. For the following treatments, fish were immersed in fish system water containing the following drugs: retinoic acid 5 µM, nocodazole 5 µM, FK506 3 µM. Fish were set out in drug solution 1 h prior injury, and solutions were changed daily. For the entire duration of the experiment, fish were kept in an incubator at 25 °C in the dark at 1 fish/100 ml density in fish system water. Negative control groups were injected or soaked, respectively, with the corresponding DMSO concentration.

### Whole mount immunohistochemistry

Fins were fixed overnight with 4% PFA at 4°C. After 2 x 5 min washes with PBTx (1x PBS with 0.5% TritonX 100) at RT, fins were transferred to 100% acetone, rinsed once and kept for 3 h at −20°C. Fins were transferred to PBTx, washed 2 x 5 min, 2 x 15 min and incubated for 30 min at RT, followed by blocking in 1% BSA in PBTx for at least 1 h at RT. Primary antibodies were diluted to 1:300 in blocking solution and fins were incubated overnight at 4°C. The next day, fins were washed several times with PBTx and incubated with secondary antibodies at 1:300 dilution in PBTx overnight at 4°C. Fins were mounted in Vectashield (Vectorlabs, H-1000). For anti-Runx2 staining, fins were fixed with 80% methanol / 20% DMSO overnight at 4°C and rehydrated by a graded series of methanol in PBTx (75, 50, 25%) for 5 min each. Fins were washed in PBTx for 30 min, after which fins were processed following the above-mentioned protocol. Primary antibodies used in this study were: mouse anti-Zns5 (Zebrafish International Resource Center, Eugene, OR, USA, RRID:AB_10013796), mouse anti-Runx (Santa Cruz sc-101145, RRID:AB_1128251), and rabbit anti-Osterix (Santa Cruz sc- 22536-R, RRID:AB_831618).

### RNAscope whole mount in situ hybridization

For detection of RNA transcripts in whole mount fins, the RNAscope Multiplex Fluorescent Reagent Kit v2 (ACD Bio-techne, 323100) was used with the following probes: *bglap*-C1 (ACD Bio-techne, 519671) and *cyp26b1*-C2 (ACD Bio-techne, 571281). Fins were fixed overnight in 4% PFA at 4°C. The next day, fins were washed with PBT (1x PBS with 0.1% Tween) 3 x 5 min each. After incubation in RNAscope Hydrogen Peroxide for 10 min, fins were rinsed 3 x with PBT, treated with RNAscope Protease Plus for 20 min, and rinsed 3 x with PBT. All these steps were performed at RT. Probes and probe diluent were pre-warmed to 40°C and cooled down to RT before use. The fins were incubated with probes overnight at 40°C. The next day, fins were washed 3x with 0.2x SSCT, 12 min each at RT. All the steps mentioned from here on were followed by such washing steps. Fins were post fixed with 4% PFA for 10 min at RT and incubated with AMP1, AMP2 and AMP3 for 30, 15 and 30 min, respectively, at 40°C. To develop the signal, fins were incubated with the corresponding RNAscope Multiplex FL v2 HRP for 15 min at 40°C. TSA-fluorophore (Perkin Elmer NEL701A001KT) was used in a 1:1500 dilution in TSA buffer (RNAscope kit). Fins were incubated for 15 min at 40°C.

### Proliferation assay

To analyse cell proliferation in the hemiray removal assay, 5-ethynyl-2′-deoxyuridine (EdU)- Click 647 kit (baseclick GmbH BCK-EdU647) was used. Fish were intraperitoneally injected with 10 µl of 10 mM EdU in PBS using a 100 µl Nanofil syringe (World Precision Instruments NANOFIL-100) once at 1 dpi for evaluation at 2 dpi, and at 1 and 2 dpi for evaluation at 3 dpi. Fins were fixed with 4% PFA overnight and AB stained following the immunohistochemistry protocol. For proliferation analysis after PMX205 treatment, fish were injected at 1 dpa and 6 h prior to harvest.

### Apoptosis assay

To analyse apoptosis, the ApopTag Red In Situ Apoptosis Detection Kit (Merck, S7165) was used. Fins were fixed overnight in 80% Methanol / 20% DMSO at 4°C. The next day, fins were rehydrated through a Methanol/PBTx (PBS + 0.5% TritonX 100) series (75%, 50%, 25%; 5 min each) and permeabilized with acetone for 3 h at −20°C. Fins were transferred to PBTx, washed 2 x 5 min, and incubated for 30 min at RT, followed by blocking in 1% BSA in PBTx for at least 1 h at RT. Mouse anti-Zns5 (Zebrafish International Resource Center, Eugene, OR, USA, RRID:AB_10013796) was diluted to 1:300 in blocking solution and fins were incubated overnight at 4°C. The next day, fins were washed several times with PBTx and transferred to superfrost slides. Using a hydrophobic pen, a circle was drawn around the fins. Fins were first equilibrated with 75 µl equilibration buffer (ApopTag kit) for 10 sec at RT, followed by incubation in 55 µl TdT enzyme (ApopTag kit) in a humidified chamber at 37°C for 1 h. The reaction was stopped by several washes with Stop/Wash buffer (ApopTag kit), followed by an incubation in Stop/Sash buffer for 10 min at RT. Fins were washed 3x 1 min with PBS and incubated in anti-DIG-Rhodamine diluted 1:2000 in blocking solution (ApopTag kit) and secondary antibody (1:300) overnight at 4°C. The next day, fins were washed 6x 20 min at RT and mounted with Vectashield (Vectorlabs, H-1000).

### Alizarin red S staining

In vivo staining with alizarin red S (ARS; Sigma-Aldrich A3757) was performed as previously described (Bensimon-Brito *et al*., 2016). Briefly, fish were stained in 0.01% ARS in system water for 15 min at RT, washed 3x 5 min in system water, immediately anaesthetized and imaged.

### Cre-lox recombination & heat-shocks

Adult double transgenic fish (*OlSp7*:CreERT2-p2a-mCherry^tud8^; *hsp70l*:loxP DsRed2 loxP nlsEGFP^tud9^, *OlSp7*:CreERT2-p2a-mCherry^tud8^; *hsp70l*:loxP Luc2-myc Stop loxP nYPet-p2a- IκBSR*, cryaa*:AmCyan^ulm15Tg^ and *OlSp7*:CreERT2-p2a-mCherry^tud8^; *hsp70l*:loxP Luc-myc STOP loxP IKKca-t2a-nls-mTagBFP2-V5, *cryaa*:AmCyan^ulm12Tg^) were intraperitoneally injected with 10 µl of 3.4 mM 4-OH tamoxifen (4-OHT) in 25% ethanol once daily for 4 days. Fish were heat-shocked twice (at 24 h and 3 h prior to harvest) at 37°C for 1 h after which water temperature was returned to 27°C within 15 minutes.

### Imaging

Images in Figures 1A, B, G; 2C, G; 4D; 5J and Suppl. Figures 2C, D were acquired with a Leica M205FA stereo microscope and display live fluorescence of fluorescent proteins. High resolution optical sections were obtained with a Leica SP8 confocal microscope or a Zeiss AxioObserver 7 equipped with an Apotome and processed using Fiji (Schindelin *et al*., 2012). Images in Figures 1C; 3A; 5B, G; Suppl. Figures 1A, B, E; Suppl. Figures 3C, D and movie S1 are z-projections. Figure 4A shows z-planes. Figure 1B contains an epifluorescence image obtained with a Zeiss AxioObserver 7. Movie S2 was recorded with a Leica M205FA stereo microscope. Movie annotations were added using the Annotate_movie plugin (Daetwyler, Modes and Fiolka, 2020).

### Cell morphology quantification

To quantify osteoblast cell shape and orientation, the transgenic line *bglap*:GFP in combination with Zns5 labelling was used. Using Fiji (Schindelin *et al*., 2012), the longest axis of a GFP+ Zns5+ cell was measured as maximum length, the short axis as maximum width, and the ratio calculated. Simultaneously, the angle of the maximum length towards the proximodistal ray axis was measured for angular deviation.

### Bulk migration assay

To quantify osteoblast migration, fins of *bglap*:GFP transgenic fish were imaged with a GFP filter and in brightfield at 0 and 1 dpa with a Leica Stereomicroscope M205FA. Using Fiji (Schindelin *et al*., 2012), a threshold was set for the GFP signal to exclude background fluorescence and to select the bulk of GFP+ cells. The thresholded image was merged with the corresponding brightfield image and the distance between GFP+ cells and the joint at the ventral-distal centre of the segment was measured. For each fin, the 2^nd^ and 3^rd^ dorsal ray was analyzed. Statistical analysis was performed on the difference in distance between 0 and 1 dpa, while for illustration the change at 1 dpa is plotted, with migration across the whole distance to the joint counting as 100% migration.

### RNAscope quantification

For quantification of RNAscope signals, optical sections were acquired with a Zeiss AxioObserver 7 microscope equipped with an Apotome, using identical imaging settings within one experiment. To analyse *bglap* intensity along the proximodistal axis, subsequent ROIs (segment height x 50 µm width) were analysed (joints excluded), and intensity was normalized for each ray. For spatial resolution of single osteoblast intensity, segment lengths were normalized and x-location of osteoblasts grouped into proximal (0.0 – 0.2), central (0.4 – 0.6) and distal (0.8 – 1.0).

### Quantification of GFP intensity

To calculate the change in *bglap*:GFP intensity in segment 0 after amputation, caudal fins were imaged at 0 and 3 dpa with a Leica Stereomicroscope M205FA. Using Fiji image analysis software (Schindelin *et al*., 2012), segment 0 of all fin rays was manually selected as region of interest, and the RawIntegratedDensity of the GFP signal was measured. Images were thresholded to exclude background fluorescent noise. The ratio of GFP intensity between 0 dpa and 3 dpa was calculated for each fish. For testing significance of intensity change differences between drug treatments and negative control fish, the RawIntegratedDensity data were box cox transformed using the formula 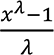 with λ = 0.2626, which we determined previously (Mishra *et al*., 2020). The difference between day 0 and day 3 was then calculated on the transformed values for each drug-treated and negative control fish, and a two-tailed student’s t- test was performed to analyse significance. For illustration, GFP intensity on day 3 is presented as % of that at day 0.

### Regenerative growth quantification

Regenerative growth was analysed in brightfield images taken at 3 dpa. For each fin, the length of the 2^nd^ and 3^rd^ dorsal ray regenerate was measured from the amputation plane to the distal tip and the average calculated. For testing significance of growth differences between drug treatments and control fish, the data were box cox transformed using the formula 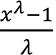 with λ = 1.5, which we determined previously (Mishra *et al*., 2020).

### Statistical analysis

Fish were randomly allocated into experimental groups. Statistical analyses were performed using GraphPad Prism (GraphPad software, LCC). For proliferation analysis in the hemiray removal model, a random distribution set was generated with MS Excel (Microsoft Corporation). Percentage data were arcsin transformed for statistical analysis. Effect size and smallest significant difference were computed to support statements about the lack of differences between experimental groups as described previously (Bertozzi *et al*., 2022). Using G*Power (Erdfelder *et al*., 2009) the effect size was computed with these parameters: t-test, tails = two, α = 0.05, power (1-β) = 0.8 and the sample size of both groups. For non-normal distributed data sets, data was log-transformed. The smallest difference that would have been significant was calculated based on the effect size and the SDs and sample sizes of the experimental groups. These smallest significant differences and the observed differences are reported in the Figure legends.

## Acknowledgements

We thank Doris Weber for her contributions to fish care and the core facility “Confocal and multiphoton imaging” of the Medical faculty of Ulm University for help with imaging. This work was funded by the Deutsche Forschungsgemeinschaft (DFG, German Research Foundation) – Project-ID 251293561 – SFB 1149, project ID 316249678 − SFB 1279, and project ID 450627322 – SFB 1506). I.S. was funded by the Hertha-Nathorff-Program of the Medical Faculty of Ulm University.

## Author contributions

Study design: I.S. and G.W. Experiments were performed and analysed by I.S. A.I., M.H.-L. and M.H.-L. provided materials. I.S. and G.W. wrote the manuscript. A.I., M.H.-L. and M.H.-L. assisted with study conceptualization and critically revised the manuscript. Funding acquisition: I.S. and G.W. All authors have read and agreed to the published version of the manuscript.

---

**Figure 1- figure supplement 1.**
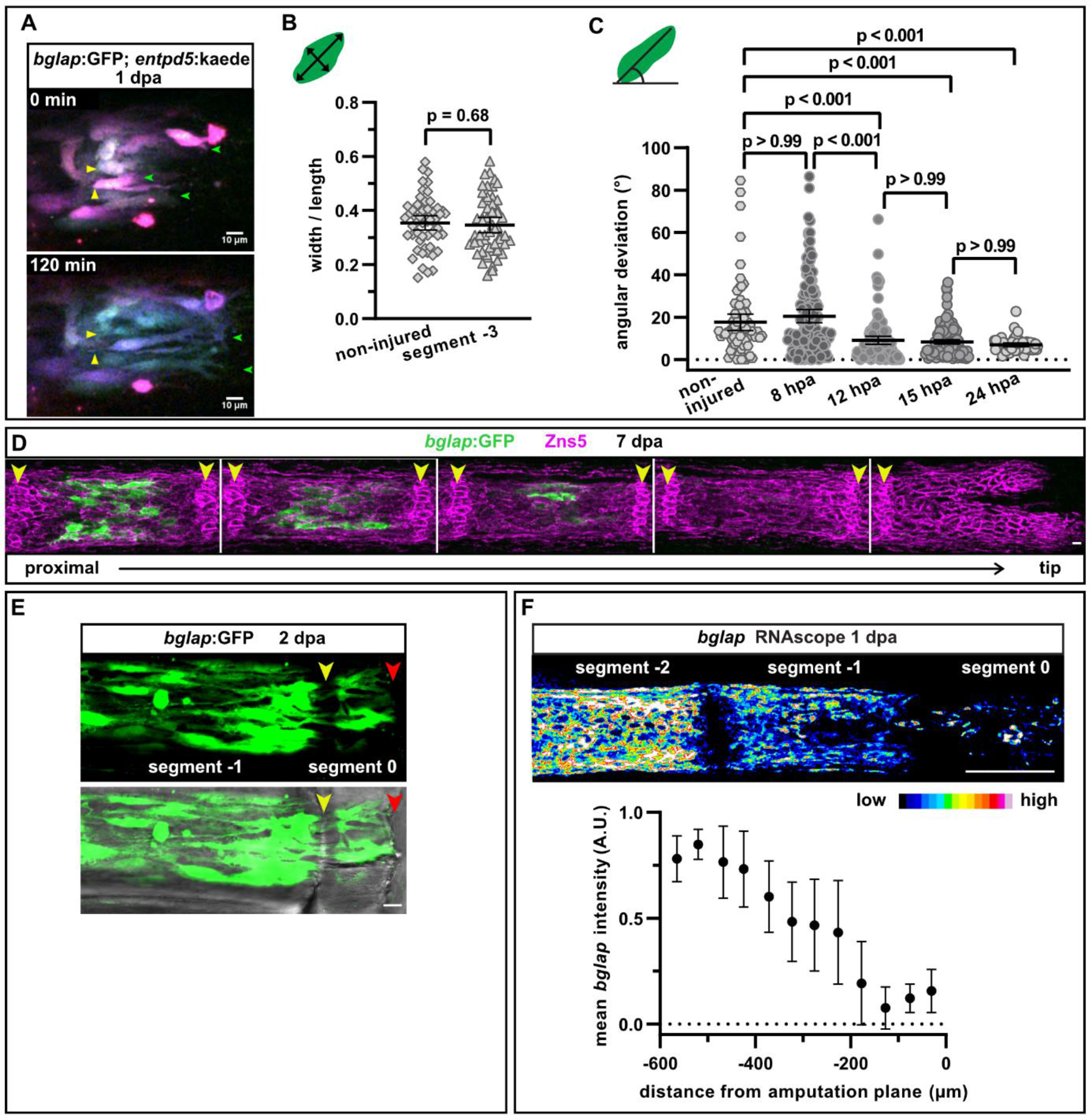
A) Still images from video 1. *bglap*:GFP; *entpd5*:kaede osteoblasts at 1 dpa. KaedeGreen was partially converted to kaedeRed, resulting in different colouring for each cell. Stationary white arrowheads point at moving cell bodies, green arrowhead at forming protrusions. Distal to the right. Scale bar, 10 µm. B) Osteoblast roundness in a segment of the non-injured fin and segment −3 at 1 dpa. *bglap*:GFP+ osteoblasts in segment −3 have a similar morphology as osteoblasts in a non-injured fin. n (fins) = 5, n (rays) = 5, n (cells) = 54. Error bars represent 95% CI. Unpaired t-test. The observed relative difference is 2%, the calculated smallest significant difference 13% and thus smaller than what we observe between segment −2 and segment −1 (Figure 1J, 54%). C) Osteoblast orientation at various time points post fin amputation. Alignment along the proximodistal axis can first be detected at 12 hpa. n (rays) = 5 (non-injured), 10 (8 hpa), 8 (12 hpa), 12 (15 hpa), 11 (24 hpa); n (cells) = 74 (non-injured), 134 (8 hpa), 124 (12 hpa), 162 (15 hpa), 53 (24 hpa). Error bars represent 95% CI. Kruskal-Wallis test. D) In the fin regenerate, the number of osteoblasts expressing the *bglap*:GFP transgene progressively increases in further proximally located (older) segments, and *bglap*:GFP+ osteoblasts are only located at the centre of each segment. The five distal segments of one fin ray are shown at 7 dpa, with osteoblasts and joint cells labelled with the pan-osteoblastic marker Zns5. In the two distal-most segments, osteoblasts are not yet mature enough to express *bglap*:GFP. Arrowheads indicate the joints. Scale bar, 10 µm. E) At 2 dpa, *bglap*:GFP+ osteoblasts can be observed within the distal joint of segment −1 and GFP+ protrusions can be seen spanning the joint, indicating that osteoblasts migrate through the joint. Yellow arrowheads, joints; red arrowheads, amputation plane. Scale bar, 10 µm. F) RNAscope in situ analysis of *bglap* expression in segments 0, −1 and −2 at 1 dpa. Expression of *bglap* gradually decreases towards the amputation plane. n (fins) = 5, n (rays) = 15. Error bars represent SEM. Scale bar, 100 µm. **Video 1. Osteoblast migration.** Live imaging of osteoblasts at 1 dpa in segment −1 (distal to the right) in a double transgenic line expressing *bglap*:GFP and *entpd5*:kaede. Partial conversion of green kaede into red kaede results in a different colouring for each cell. Stationary yellow arrowheads highlight the migration of cell bodies towards the amputation plane and retraction of the rear ends. Additionally, several cells can be observed forming long protrusions extending distally (green arrowheads). Distal to the right. Scale bar, 10 µm.

**Figure 4 – figure supplement 1.**
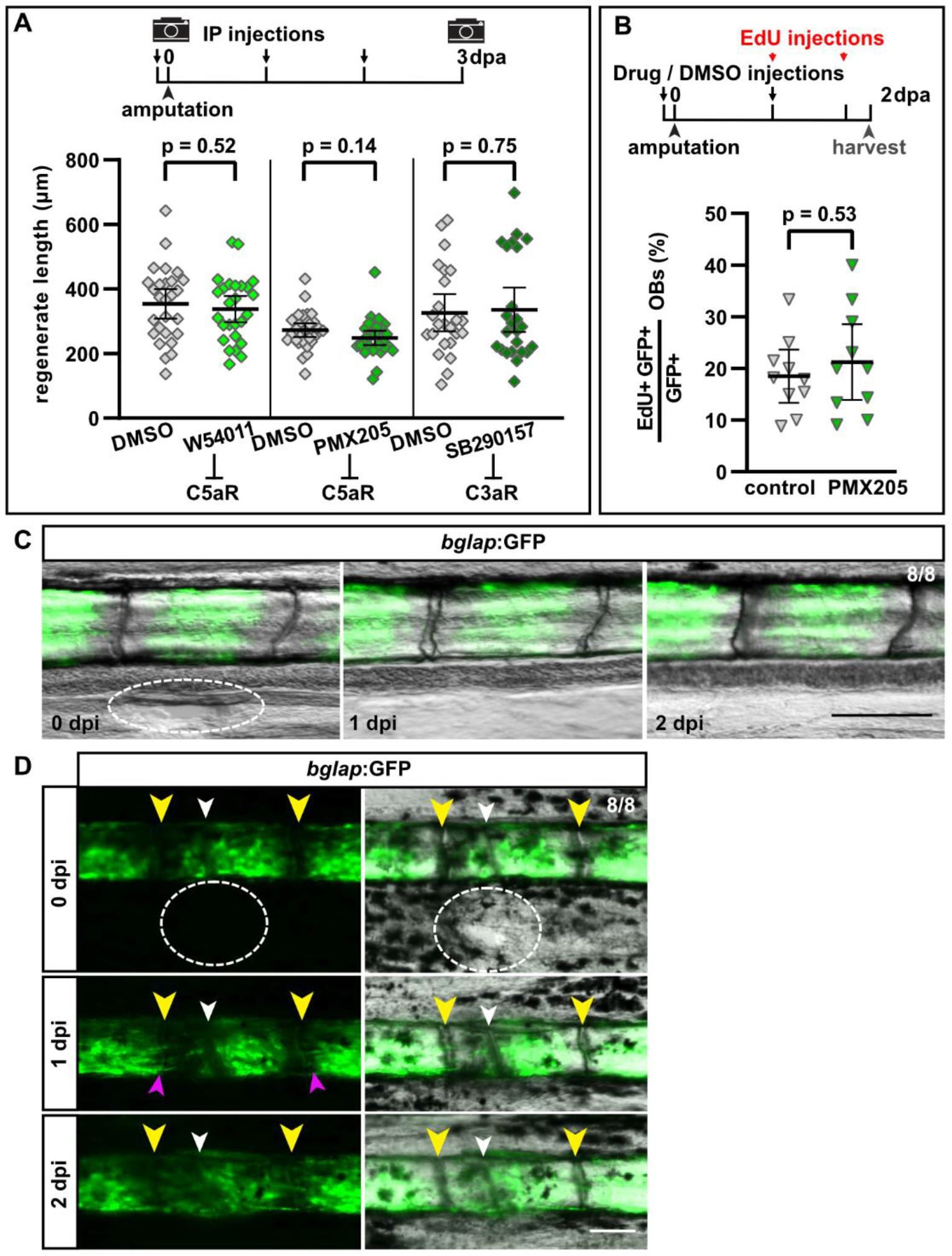
A) Absolute regenerate length at 3 dpa is not affected by W54011, PMX205 or SB290157 treatment. N (experiments) = 3, n (fins) = 27 (W54011), 29 (PMX205), 24 (SB290157); appertaining controls have the same n. Error bars represent 95% CI. Unpaired t-tests. The observed relative difference is 8% (W54011), 13% (PMX205) and 6% (SB290157), the calculated smallest significant differences are 30% (W54011), 21% (PMX205) and 36% (SB290157), which are smaller than what we have observed previously in a large chemical screen (Mishra *et al*., 2020), suggesting that in principle our experiment had enough power to detect a similar effect size. B) The fraction of proliferating osteoblasts (EdU+ *bglap*:GFP+) at 2 dpa is not altered by I.P. injection of PMX205. N (experiments) = 1, n (fish) = 10. Error bars represent 95% CI. Unpaired t-tests. The observed relative difference is 7%, the calculated smallest significant difference is 28%. C) Incision in the interray skin (dashed oval) close to the bony ray does not induce migration of *bglap*:GFP+ osteoblasts from the bony ray into the interray mesenchyme. n = 8/8 fins show no migration. Scale bar, 100 µm. D) Incision in the interray (dashed oval) in combination with bone fracture (white arrowhead) induces migration of *bglap*:GFP+ osteoblasts towards the fracture (pink arrowheads), but not into the interray. Yellow arrowheads indicate joints. n = 8/8 fins. Scale bar, 100 µm.

**Figure 5 – figure supplement 1.**
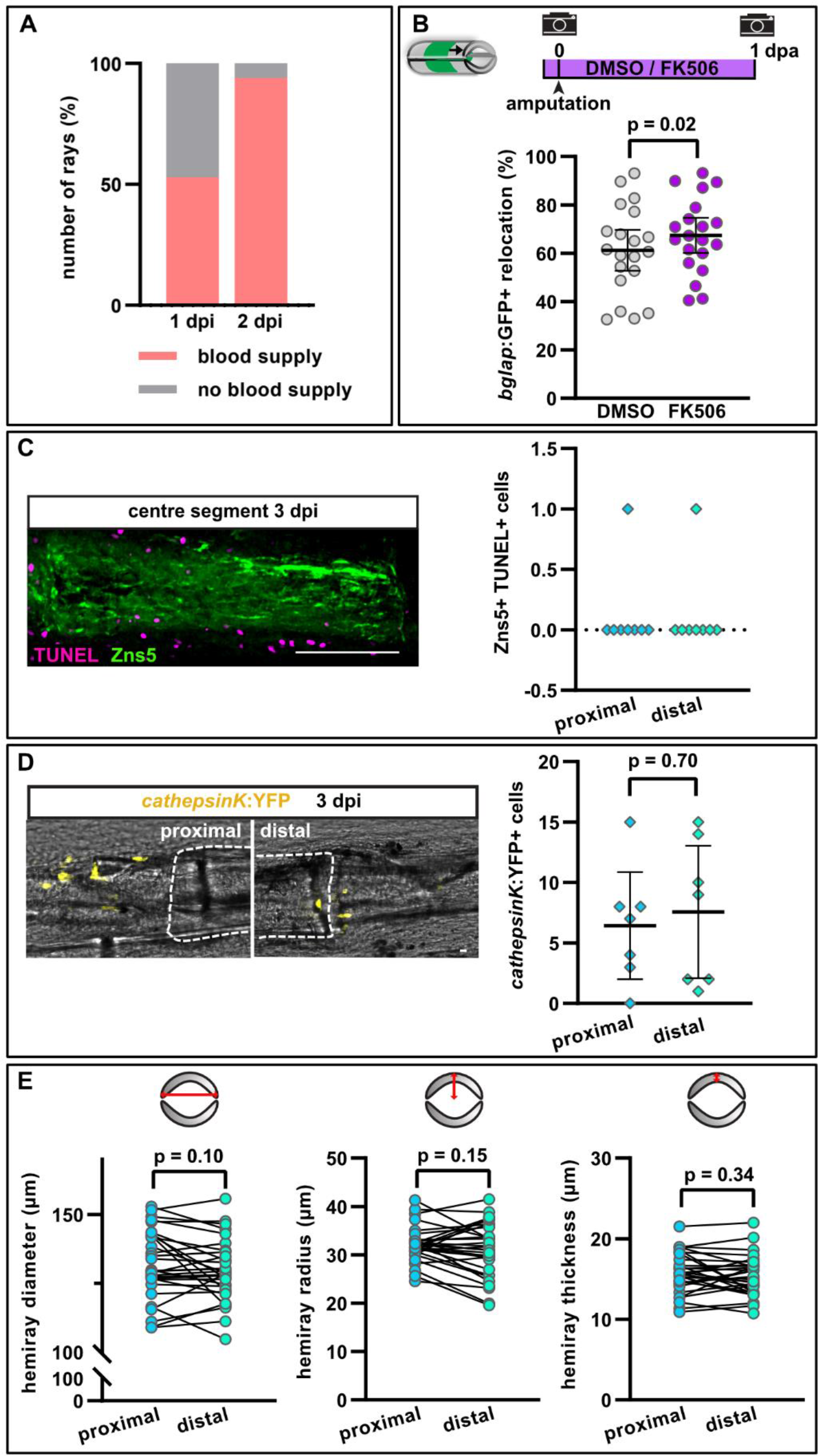
A) Re-vascularization of the centre segment at 1 and 2 dpi. Most segments are fully recovered by 2 dpi. n (segments) = 36. B) Calcineurin inhibition with FK506 slightly increases osteoblast recruitment towards the amputation plane in segment −1. 100% indicates full crossing of the distance to the respective joint at 1 dpa. N (experiments) = 1, n (fins) = 10, n (rays) = 20. Error bars represent 95% CI. Unpaired t-test. C) TUNEL staining for apoptotic cells with Zns5 labelling of osteoblasts in the centre segment at 3 dpi. n (segments) = 8. Scale bar, 100 µm. D) Number of osteoclasts labelled using the *cathepsinK*:YFP transgenic line at the centre segment at 3 dpi. No difference is detected between the proximal and distal injury site. n (segments) = 7. Error bars represent 95% CI. Unpaired t-test. Scale bar, 100 µm. E) Bone matrix features at the proximal and distal end of the 3^rd^ segment proximal to the bifurcation of the 2^nd^ dorsal ray (which is the segment used as centre segment in the hemiray removal assays). The diameter and radius of a hemiray is analysed, as well as the thickness. No difference between the proximal and distal end can be detected. n (fins) = 15, n (segments) = 30. Paired t- tests **Video 2.** Re-vascularization of the centre segment Live imaging of blood flow at 2 dpi. Yellow arrowheads indicate segment borders of the centre segment. Lower ray is not injured. Distal to the right. Scale bar, 100 µm.

## References

Aleström, P. et al. (2020) ‘Zebrafish: Housing and husbandry recommendations’, Laboratory animals, 54(3), pp. 213–224. doi: 10.1177/0023677219869037.

Aman, A. and Piotrowski, T. (2010) ‘Cell migration during morphogenesis’, Developmental Biology, 341(1), pp. 20–33. doi: 10.1016/j.ydbio.2009.11.014.

Ames, R. S. et al. (2001) ‘Identification of a Selective Nonpeptide Antagonist of the Anaphylatoxin C3a Receptor That Demonstrates Antiinflammatory Activity in Animal Models’, The Journal of Immunology, 166(10), pp. 6341–6348. doi: 10.4049/jimmunol.166.10.6341.

Ando, K. et al. (2017) ‘Osteoblast Production by Reserved Progenitor Cells in Zebrafish Bone Regeneration and Maintenance.’, Developmental cell, 43(5), pp. 643–650.e3. doi: 10.1016/j.devcel.2017.10.015.

Van Antwerp, D. et al. (1996) ‘Suppression of TNF-alpha-induced apoptosis by NF-kappaB’, Science (New York, N.Y.), 274(5288), pp. 787–789. doi: 10.1126/SCIENCE.274.5288.787.

Apschner, A. et al. (2014) ‘Zebrafish enpp1 mutants exhibit pathological mineralization, mimicking features of generalized arterial calcification of infancy (GACI) and pseudoxanthoma elasticum (PXE)’, DMM Disease Models and Mechanisms, 7(7), pp. 811–822. doi: 10.1242/DMM.015693/-/DC1.

Arbore, G. et al. (2016) ‘T helper 1 immunity requires complement-driven NLRP3 inflammasome activity in CD4^+^ T cells’, Science (New York, N.Y.), 352(6292). doi: 10.1126/SCIENCE.AAD1210.

Arbore, G., Kemper, C. and Kolev, M. (2017) ‘Intracellular complement − the complosome − in immune cell regulation’, Molecular Immunology. doi: 10.1016/j.molimm.2017.05.012.

Bensimon-Brito, A. et al. (2016) ‘Revisiting in vivo staining with alizarin red S--a valuable approach to analyse zebrafish skeletal mineralization during development and regeneration’, BMC developmental biology, 16(1). doi: 10.1186/S12861-016-0102-4.

Bertozzi, A. et al. (2022) ‘Wnt/β-catenin signaling acts cell-autonomously to promote cardiomyocyte regeneration in the zebrafish heart’, Developmental biology, 481, pp. 226–237. doi: 10.1016/J.YDBIO.2021.11.001.

Blum, N. and Begemann, G. (2015) ‘Osteoblast de- and redifferentiation are controlled by a dynamic response to retinoic acid during zebrafish fin regeneration’, Development. 2015/08/09, 142(17), pp. 2894–2903. doi: 10.1242/dev.120204.

Böhm, A.-M. et al. (2019) ‘Activation of Skeletal Stem and Progenitor Cells for Bone Regeneration Is Driven by PDGFRβ Signaling’, Developmental Cell, 51(2), pp. 236–254.e12. doi: 10.1016/j.devcel.2019.08.013.

Boshra, H., Li, J. and Sunyer, J. O. (2006) ‘Recent advances on the complement system of teleost fish’, Fish & Shellfish Immunology, 20(2), pp. 239–262. doi: 10.1016/J.FSI.2005.04.004.

Brown, S. S. and Spudich, J. A. (1979) ‘Cytochalasin inhibits the rate of elongation of actin filament fragments’, Journal of Cell Biology, 83(3), pp. 657–662. doi: 10.1083/jcb.83.3.657.

Brown, S. S. and Spudich, J. A. (1981) ‘Mechanism of action of cytochalasin: evidence that it binds to actin filament ends’, Journal of Cell Biology, 88(3), pp. 487–491. doi: 10.1083/jcb.88.3.487.

Cao, Z. et al. (2021) ‘Calcineurin controls proximodistal blastema polarity in zebrafish fin regeneration’, Proceedings of the National Academy of Sciences, 118(2), p. e2009539118. doi: 10.1073/pnas.2009539118.

Chablais, F. and Jaźwińska, A. (2010) ‘IGF signaling between blastema and wound epidermis is required for fin regeneration’, Development, 137(6), pp. 871–879. doi: 10.1242/dev.043885.

Chen, C. H. et al. (2015) ‘Transient laminin beta 1a Induction Defines the Wound Epidermis during Zebrafish Fin Regeneration’, PLoS Genet. 2015/08/26, 11(8), p. e1005437. doi: 10.1371/journal.pgen.1005437.

Chugh, P. and Paluch, E. K. (2018) ‘The actin cortex at a glance’, Journal of Cell Science, 131(14). doi: 10.1242/jcs.186254.

Daetwyler, S., Modes, C. D. and Fiolka, R. (2020) ‘Fiji plugin for annotating movies with custom arrows’, Biology Open, 9(11). doi: 10.1242/BIO.056200/266550/AM/FIJI-PLUGIN-FOR-ANNOTATING-MOVIES-WITH-CUSTOM.

Daveau, M. et al. (2004) ‘Expression of a functional C5a receptor in regenerating hepatocytes and its involvement in a proliferative signaling pathway in rat’, Journal of immunology (Baltimore, Md. : 1950), 173(5), pp. 3418–3424. doi: 10.4049/JIMMUNOL.173.5.3418.

Dirckx, N., Van Hul, M. and Maes, C. (2013) ‘Osteoblast recruitment to sites of bone formation in skeletal development, homeostasis, and regeneration’, Birth defects research. Part C, Embryo today : reviews, 99(3), pp. 170–191. doi: 10.1002/BDRC.21047.

Ehrnthaller, C. et al. (2013) ‘Complement C3 and C5 deficiency affects fracture healing’, PLoS One. 2013/11/22, 8(11), p. e81341. doi: 10.1371/journal.pone.0081341.

Erdfelder, E. et al. (2009) ‘Statistical power analyses using G*Power 3.1: tests for correlation and regression analyses’, Behavior research methods, 41(4), pp. 1149–1160. doi: 10.3758/BRM.41.4.1149.

Espinosa, A. V. et al. (2009) ‘Regulator of calcineurin 1 modulates cancer cell migration in vitro’, Clinical and Experimental Metastasis, 26(6), pp. 517–526. doi: 10.1007/S10585-009-9251-1/FIGURES/5.

Etienne-Manneville, S. (2013) ‘Microtubules in Cell Migration’, Annual Review of Cell and Developmental Biology, 29(1), pp. 471–499. doi: 10.1146/annurev-cellbio-101011-155711.

Florian, S. and Mitchison, T. J. (2016) ‘Anti-microtubule drugs’, in *Methods in Molecular Biology*. Humana Press Inc., pp. 403–421. doi: 10.1007/978-1-4939-3542-0_25.

Gemberling, M. et al. (2013) ‘The zebrafish as a model for complex tissue regeneration’, Trends Genet. 2013/08/10, 29(11), pp. 611–620. doi: 10.1016/j.tig.2013.07.003.

Geurtzen, K. et al. (2014) ‘Mature osteoblasts dedifferentiate in response to traumatic bone injury in the zebrafish fin and skull’, Development. 2014/05/14, 141(11), pp. 2225–2234. doi: 10.1242/dev.105817.

Huber-Lang, M., Kovtun, A. and Ignatius, A. (2013) ‘The role of complement in trauma and fracture healing’, Semin Immunol. 2013/06/19, 25(1), pp. 73–78. doi: 10.1016/j.smim.2013.05.006.

Huitema, L. F. A. et al. (2012) ‘Entpd5 is essential for skeletal mineralization and regulates phosphate homeostasis in zebrafish.’, Proceedings of the National Academy of Sciences of the United States of America, 109(52), pp. 21372–7. doi: 10.1073/pnas.1214231110.

Ignatius, A., Schoengraf, P., et al. (2011) ‘Complement C3a and C5a modulate osteoclast formation and inflammatory response of osteoblasts in synergism with IL-1β’, Journal of Cellular Biochemistry, 112(9), pp. 2594–2605. doi: 10.1002/JCB.23186.

Ignatius, A., Ehrnthaller, C., et al. (2011) ‘The anaphylatoxin receptor C5aR is present during fracture healing in rats and mediates osteoblast migration in vitro.’, The Journal of trauma, 71(4), pp. 952–60. doi: 10.1097/TA.0b013e3181f8aa2d.

Jain, U., Woodruff, T. and Stadnyk, A. (2013) ‘The C5a receptor antagonist PMX205 ameliorates experimentally induced colitis associated with increased IL-4 and IL-10’, British Journal of Pharmacology, 168(2), pp. 488–501. doi: 10.1111/j.1476-5381.2012.02183.x.

Jung, H. and Oh, E. S. (2016) ‘FK506 positively regulates the migratory potential of melanocyte-derived cells by enhancing syndecan-2 expression’, Pigment cell & melanoma research, 29(4), pp. 434–443. doi: 10.1111/PCMR.12480.

Kimura, Y. et al. (2003) ‘Expression of Complement 3 and Complement 5 in Newt Limb and Lens Regeneration’, The Journal of Immunology, 170(5), pp. 2331–2339. doi: 10.4049/jimmunol.170.5.2331.

Knopf, F. et al. (2011) ‘Bone regenerates via dedifferentiation of osteoblasts in the zebrafish fin’, Dev Cell. 2011/05/17, 20(5), pp. 713–724. doi: 10.1016/j.devcel.2011.04.014.

Kovács, M. et al. (2004) ‘Mechanism of blebbistatin inhibition of myosin II’, Journal of Biological Chemistry, 279(34), pp. 35557–35563. doi: 10.1074/jbc.M405319200.

Kujawski, S. et al. (2014) ‘Calcineurin regulates coordinated outgrowth of zebrafish regenerating fins.’, Developmental cell, 28(5), pp. 573–87. doi: 10.1016/j.devcel.2014.01.019.

Kular, J. et al. (2012) ‘An overview of the regulation of bone remodelling at the cellular level’, Clinical biochemistry, 45(12), pp. 863–873. doi: 10.1016/J.CLINBIOCHEM.2012.03.021.

Liszewski, M. K. et al. (2013) ‘Intracellular Complement Activation Sustains T Cell Homeostasis and Mediates Effector Differentiation’, Immunity, 39(6), pp. 1143–1157. doi: 10.1016/J.IMMUNI.2013.10.018.

Liu, Y. et al. (2010) ‘Calcineurin promotes proliferation, migration, and invasion of small cell lung cancer’, Tumor Biology, 31(3), pp. 199–207. doi: 10.1007/S13277-010-0031-Y/FIGURES/5.

March, D. R. et al. (2004) ‘Potent Cyclic Antagonists of the Complement C5a Receptor on Human Polymorphonuclear Leukocytes. Relationships between Structures and Activity’, Molecular Pharmacology, 65(4), pp. 868–879. doi: 10.1124/mol.65.4.868.

Marí-Beffa, M. and Murciano, C. (2010) ‘Dermoskeleton morphogenesis in zebrafish fins’, Developmental Dynamics, 239(11), pp. 2779–2794. doi: 10.1002/dvdy.22444.

Markiewski, M. M. et al. (2004) ‘C3a and C3b activation products of the third component of complement (C3) are critical for normal liver recovery after toxic injury’, Journal of immunology (Baltimore, Md. : 1950), 173(2), pp. 747–754. doi: 10.4049/JIMMUNOL.173.2.747.

Mastellos, D. et al. (2001) ‘A novel role of complement: mice deficient in the fifth component of complement (C5) exhibit impaired liver regeneration’, Journal of immunology (Baltimore, Md. : 1950), 166(4), pp. 2479–2486. doi: 10.4049/JIMMUNOL.166.4.2479.

Mastellos, D. and Lambris, J. D. (2002) ‘Complement: more than a “guard” against invading pathogens?’, Trends in immunology, 23(10), pp. 485–491. doi: 10.1016/S1471-4906(02)02287-1.

Merle, N. S. et al. (2015) ‘Complement system part I − molecular mechanisms of activation and regulation’, Frontiers in Immunology, 6(JUN), p. 262. doi: 10.3389/FIMMU.2015.00262/BIBTEX.

Mierke, C. T. (2015) ‘Physical view on migration modes’, Cell Adh Migr. 2015/07/21, 9(5), pp. 367–379. doi: 10.1080/19336918.2015.1066958.

Mishra, R. et al. (2020) ‘NF-κB Signaling Negatively Regulates Osteoblast Dedifferentiation during Zebrafish Bone Regeneration’, Developmental Cell, 52(2), pp. 167–182.e7. doi: 10.1016/j.devcel.2019.11.016.

Miyajima, D. et al. (2012) ‘Profilin1 Regulates Sternum Development and Endochondral Bone Formation’, Journal of Biological Chemistry, 287(40), pp. 33545–33553. doi: 10.1074/JBC.M111.329938.

Natarajan, N. et al. (2018) ‘Complement receptor C5AR1 plays an evolutionarily conserved role in successful cardiac regeneration’, Circulation, 137(20), pp. 2152–2165. doi: 10.1161/CIRCULATIONAHA.117.030801.

Owlarn, S. et al. (2017) ‘Generic wound signals initiate regeneration in missing-tissue contexts’, Nature Communications, 8(1), p. 2282. doi: 10.1038/s41467-017-02338-x.

Petrie, R. J., Doyle, A. D. and Yamada, K. M. (2009) ‘Random versus directionally persistent cell migration.’, Nature reviews. Molecular cell biology, 10(8), pp. 538–49. doi: 10.1038/nrm2729.

Pfefferli, C. and Jazwinska, A. (2015) ‘The art of fin regeneration in zebrafish’, Regeneration (Oxf). 2015/04/01, 2(2), pp. 72–83. doi: 10.1002/reg2.33.

Poss, K. D. et al. (2004) ‘Germ cell aneuploidy in zebrafish with mutations in the mitotic checkpoint gene mps1’, Genes & Development, 18(13), pp. 1527–1532. doi: 10.1101/GAD.1182604.

Prideaux, M. et al. (2015) ‘Parathyroid Hormone Induces Bone Cell Motility and Loss of Mature Osteocyte Phenotype through L-Calcium Channel Dependent and Independent Mechanisms’, PloS one, 10(5). doi: 10.1371/JOURNAL.PONE.0125731.

Del Rio-Tsonis, K. et al. (1998) ‘Expression of the third component of complement, C3, in regenerating limb blastema cells of urodeles.’, Journal of immunology (Baltimore, Md. : 1950), 161(12), pp. 6819–24. Available at: http://www.ncbi.nlm.nih.gov/pubmed/9862713 (Accessed: 14 April 2020).

Schindelin, J. et al. (2012) ‘Fiji: an open-source platform for biological-image analysis’, Nature methods, 9(7), pp. 676–682. doi: 10.1038/NMETH.2019.

Singh, S. P., Holdway, J. E. and Poss, K. D. (2012) ‘Regeneration of Amputated Zebrafish Fin Rays from De Novo Osteoblasts’, Developmental Cell, 22(4), pp. 879–886. doi: 10.1016/J.DEVCEL.2012.03.006.

Sousa, S. et al. (2011) ‘Differentiated skeletal cells contribute to blastema formation during zebrafish fin regeneration.’, Development (Cambridge, England), 138(18), pp. 3897–905. doi: 10.1242/dev.064717.

Stewart, S. and Stankunas, K. (2012) ‘Limited dedifferentiation provides replacement tissue during zebrafish fin regeneration’, Developmental Biology, 365(2), pp. 339–349. doi: 10.1016/J.YDBIO.2012.02.031.

Su, P. et al. (2020) ‘MACF1 promotes preosteoblast migration by mediating focal adhesion turnover through EB1.’, Biology Open, 9(3). doi: 10.1242/BIO.048173.

Su, P. et al. (2022) ‘MACF1 promotes osteoblastic cell migration by regulating MAP1B through the GSK3beta/TCF7 pathway’, Bone, 154, p. 116238. doi: 10.1016/J.BONE.2021.116238.

Sumichika, H. et al. (2002) ‘Identification of a potent and orally active non-peptide C5a receptor antagonist’, Journal of Biological Chemistry, 277(51), pp. 49403–49407. doi: 10.1074/jbc.M209672200.

Sun, L. et al. (2005) ‘Calcineurin regulates bone formation by the osteoblast’, Proceedings of the National Academy of Sciences, 102(47), pp. 17130–17135. doi: 10.1073/PNAS.0508480102.

Svitkina, T. (2018) ‘The actin cytoskeleton and actin-based motility’, Cold Spring Harbor Perspectives in Biology, 10(1). doi: 10.1101/cshperspect.a018267.

Swaney, K. F., Huang, C.-H. and Devreotes, P. N. (2010) ‘Eukaryotic Chemotaxis: A Network of Signaling Pathways Controls Motility, Directional Sensing, and Polarity’, Annual Review of Biophysics, 39(1), pp. 265–289. doi: 10.1146/annurev.biophys.093008.131228.

Tang, Y. et al. (2009) ‘TGF-beta1-induced migration of bone mesenchymal stem cells couples bone resorption with formation’, Nature medicine, 15(7), pp. 757–765. doi: 10.1038/NM.1979.

Thiel, A. et al. (2018) ‘Osteoblast migration in vertebrate bone’, Biological Reviews, 93(1), pp. 350–363. doi: 10.1111/brv.12345.

Thorgersen, E. B. et al. (2019) ‘The Role of Complement in Liver Injury, Regeneration, and Transplantation’, Hepatology. John Wiley and Sons Inc., pp. 725–736. doi: 10.1002/hep.30508.

Torreggiani, E. et al. (2013) ‘Preosteocytes/Osteocytes Have the Potential to Dedifferentiate Becoming a Source of Osteoblasts’, PLOS ONE, 8(9), p. e75204. doi: 10.1371/JOURNAL.PONE.0075204.

Vanoevelen, J. et al. (2011) ‘Trpv5/6 is vital for epithelial calcium uptake and bone formation’, The FASEB Journal, 25(9), pp. 3197–3207. doi: 10.1096/FJ.11-183145.

Wurtzel, O. et al. (2015) ‘A Generic and Cell-Type-Specific Wound Response Precedes Regeneration in Planarians In Brief Developmental Cell Resource A Generic and Cell-Type- Specific Wound Response Precedes Regeneration in Planarians’. doi: 10.1016/j.devcel.2015.11.004.

Zhang, S. and Cui, P. (2014) ‘Complement system in zebrafish’, Developmental & Comparative Immunology. 2014/01/28, 46(1), pp. 3–10. doi: 10.1016/j.dci.2014.01.010.

Zhang, W. et al. (2014) ‘VEGF and BMP-2 promote bone regeneration by facilitating bone marrow stem cell homing and differentiation’, European cells & materials, 27, pp. 1–12. doi: 10.22203/ECM.V027A01.

Zhang, X. et al. (2011) ‘Rho kinase inhibitors stimulate the migration of human cultured osteoblastic cells by regulating actomyosin activity’, Cellular and Molecular Biology Letters, 16(2), pp. 279–295. doi: 10.2478/S11658-011-0006-Z/MACHINEREADABLECITATION/RIS.

